# Transient receptor potential vanilloid channel 2 contributes to multi-modal endoplasmic reticulum and perinuclear space dilations that can also be observed in prion-infected mice

**DOI:** 10.1101/2024.12.18.629254

**Authors:** Wenda Zhao, Shehab Eid, Chris Sackmann, Declan Williams, Xinzhu Wang, Yunqing Ouyang, Thomas Zerbes, Gerold Schmitt-Ulms

## Abstract

Our recent work on the prion protein and Na^+^,K^+^-ATPases (NKAs) led us to revisit data from over 50 years ago, which suggested a similarity between vacuolation phenotypes in rodents poisoned with cardiac glycosides (CGs) and spongiform degeneration in prion disease. At that time, this hypothesis was dismissed because the vacuolation observed in prion diseases affects neurons, whereas CG poisoning in rodent brains led to swellings of the endoplasmic reticulum (ER) in astrocytes. We speculated that this difference might be specific to rodents and document here that the vacuolation shifts to neurons in mice expressing a humanized NKA α1 subunit. Next, we investigated the molecular mechanisms that could cause similar ER vacuolation in human cells *in vitro*. We found that certain stressors—such as overexpression of NKA α subunits and exposure to specific toxins known to trigger the unfolded protein response—can induce a phenotype characterized by profound ER dilation that is most strikingly observed for the perinuclear space (PNS). The ion imbalance typically caused by functional NKAs does not contribute to this phenotype. In fact, it can occur even with the overexpression of catalytically inactive NKAs. Several lines of evidence, generated with pharmacological agents, ion-specific dyes, antagonists, and truncated expression constructs, suggest a calcium leak channel in the ER, known as transient receptor potential vanilloid 2 (TRPV2), plays a role in this ER and PNS dilation. Additionally, we observed that the formation of these vacuoles coincides with a decrease in steady-state levels of the lipid kinase PIKFYVE, which is recognized for its role in endolysosomal fission and fusion processes. Finally, we found evidence of vacuoles in cryo-sectioned brains of prion-infected mice that can be filled with a fluorescent marker targeted to the ER and PNS. This raises the possibility that this vacuolation phenomenon contributes to spongiform degeneration seen in prion diseases.

## INTRODUCTION

First reported in 1898 in a sheep afflicted with scrapie disease ^1^, a phenotype of subcellular neuronal vacuoles—today referred to as spongiform degeneration—is a defining characteristic of a group of rapidly progressive and fatal neurodegenerative diseases. In addition to scrapie in sheep, these include Creutzfeldt-Jakob disease (CJD) in humans, bovine spongiform encephalopathy (BSE) in cattle, and chronic wasting disease (CWD) in deer and elk. In 1982, a then unknown protein, termed the prion protein, was proposed as the causative agent of these diseases, now increasingly referred to as prion diseases ^2^. This discovery launched a productive line of research revealing that prion diseases occur when abnormal conformers of the prion protein, termed PrP Scrapie (PrP^Sc^), accumulate in the brains of affected animals or individuals, thereby inducing the normal cellular prion protein (PrP^C^) to convert to PrP^Sc^ through a mechanism of templated conversion ^3^.

Remarkably, although interest in the molecular underpinnings of these diseases remains high, the identity of the membrane-bound neuronal vacuoles that give rise to spongiform degeneration has not been unequivocally determined. Recent research has pointed toward the possibility that these vacuoles may have endolysosomal origins, based on observations that levels of the phosphoinositide kinase PIKFYVE were depleted in prion disease models ^4^. This conclusion was built on previous data, which established that the functional impairment of a complex composed of PIKFYVE, FIG4, and VAC14 can result in neuronal vacuoles through dilation of endolysosomal compartments ^5–7^. Although suggestive of spongiform degeneration in prion diseases sharing endolysosomal origins, alternative explanations need to be considered. For instance, a lowering of steady-state PIKFYVE levels may result from insults to the endoplasmic reticulum (ER), which can be expected to activate the unfolded protein response (UPR). One arm of this response attenuates protein translation through phosphorylation of the α- subunit of eukaryotic initiation factor 2 (eIF2α-P). This does not have to affect protein synthesis of all gene products equally, because certain stresses are understood to activate a branch of this response centered on the transcription factors ATF4 and CCAAT/enhancer-binding protein homologous protein (CHOP) that can drive selective protein translation ^8^.

Through protein-protein interactions studies of *in vivo* crosslinked brain tissue, we have learned that PrP^C^ resides in the brain in immediate proximity to Na^+^,K^+^-ATPases (NKAs) ^9^. NKAs are minimally composed of one α-subunit and one β-subunit ^10–12^. Like PrP^C^, NKAs localize primarily to raft domains within the plasma membrane ^13, 14^, where they antiport during one ATP-driven pump cycle two potassium and three sodium ions in and out of the cell, respectively ^15^. It has been proposed that in an electrically active brain cell, a majority of the cell’s total ATP may be consumed by the combined activities of NKAs as they generate the electrochemical gradient necessary for secondary transport processes ^16^. Exposure to toxic levels of cardiac glycosides (CGs) inhibits NKAs ^15, 17^. This inhibition disrupts the electrochemical gradient ^12^, leading to the collapse of critical transmembrane transport processes required to sustain cellular homeostasis.

A hypothesis that NKA inhibition might drive vacuolation in prion diseases was initially proposed in 1966 ^18^ and refined in later years ^19–22^. The hypothesis did not gain traction because the CG-induced vacuolation primarily caused swelling in astrocytes ^23, 24^, whereas spongiform degeneration in prion diseases is mainly observed in neurons ^1, 25–28^.

Intrigued by these data, we wondered if differences in relative CG susceptibilities of brain cell types in different species can account for them: Like the brains of humans, mouse brains express three different NKA α subunits, termed α1, α2, and α3 that are coded by *Atp1a1*, *Atp1a2*, and *Atp1a3* genes, respectively. Astrocytes primarily express pump molecules that include the highly CG-sensitive α2 subunit ^29, 30^. Therefore, when exposed to CGs, these cells are expected to experience a significant blockade of their sodium and potassium transport. In contrast, NKAs in neurons are made up of α1 or α3 subunits, and the α1 subunit in rodents differs from other mammalian α1 orthologs in two amino acid positions that render the rodent isoform refractory to CG inhibition ^31, 32^.

Here, we show that vacuoles can indeed also form in brain neurons of mice once their NKA α1 subunit has been sensitized to CGs through *Atp1a1* gene edits. It has previously been shown that these vacuoles represent dilations of the ER and the PNS, the latter representing the largest ER lumen in cells ^33^. This finding ties them to a growing list of insults that can cause ER and PNS dilations in a subset of cells ^34^. During this study, we used some of these paradigms and observed that the dilation phenotype was paralleled by and dependent on an increase in ER calcium levels. Based on these findings and the multi-modal nature of conditions that have been reported to induce this phenotype, we suspected a specific calcium ion leak channel in their formation. Indeed, we show that manipulations of this leak channel augment or block ER and PNS vacuolation. We also establish that these dilations are reversible if counteracted early enough and, intriguingly, trigger the UPR, leading to PIKFYVE depletion. Finally, we document that the intracerebral infection of mice with prions gives rise to PNS dilations, not observed in healthy control mice. These PNS dilations can be filled with the artificial spEGFP^KDEL^ expression construct, a well-known ER marker, which we delivered to the brain through intravenous injection of a recombinant adeno-associated virus (rAAV) vector. Based on these findings, we are proposing that PNS dilations contribute to the spongiform degeneration observed in prion diseases.

## Materials and Methods

### Chemicals

Ouabain (catalog number O3125-250MG), strophanthidin (catalog number S6626-250MG), tranilast (catalog number T0318-10MG), and 4,4′-diisothiocyanatostilbene-2,2′-disulfonic acid disodium salt hydrate (DIDS) (catalog number D3514-100MG) were purchased from MilliporeSigma (Burlington, MA, USA). Fluo-3-AM (catalog number F1241), Mag-Fluo-4-AM (catalog number M14206), BAPTA-AM (catalog number B1205), and RNase A (catalog number EN0531) were purchased from Thermo Fisher Scientific (Waltham, MA, USA). Ophiobolin A (catalog number 15381-1) and cyclosporin A (catalog number 12088-25) were purchased from Cedarlane Labs (Burlington, ON, Canada). N-(furan-2-ylmethyl)-3-[4-[methyl(propyl)amino]-6- (trifluoromethyl)pyrimidin-2-yl]sulfanylpropanamide (SET2) was purchased from MedChem Express (catalog number 2313525-20-9, Monmouth Junction, NJ, USA). Doxycycline hydrochloride (catalog number DB0889-25G) was purchased from BioBasic (Markham, ON, Canada). Hoechst 33342 (catalog number 4082S) was purchased from Cell Signaling Technology (Danvers, MA, USA). Hematoxylin and Eosin staining kit (catalog number ab245880) was purchased from Abcam (Cambridge, UK).

#### Reconstitution of chemicals

The following compounds were reconstituted in dimethylsulfoxide (DMSO) (catalog number 12611P, Cell Signaling Technology) at the indicated stock concentrations: tranilast (30 mM), DIDS (20 mM), Fluo-3-AM (1 mM), BAPTA-AM (10 mM), ophiobolin A (1 mM), cyclosporin A (20 mM), SET2 (10 mM). Other compounds were reconstituted in UltraPure water (catalog number 10977-015, Invitrogen, Waltham, MA, USA) at the indicated stock concentration: ouabain (1 mM), strophanthidin (1 mM), doxycycline hydrochloride (1 mg/mL), Hoechst 33342 (10 mg/mL).

### Antibodies

#### Primary antibodies

rabbit monoclonal IgG anti-NeuN antibody, clone EPR12763, dilution 1:1,000 (catalog number ab177487, Abcam, Cambridge, UK), rabbit monoclonal IgG anti-KDEL antibody, clone EPR12668, dilution 1:2,000 (catalog number ab176333, Abcam), mouse monoclonal IgG1 anti-ATP1A1 antibody, clone 464.6, dilution 1:2,000 (catalog number ab7671, Abcam), rabbit polyclonal anti-ATP1A2 (epitope: second cytoplasmic domain) antibody, dilution 1:2,000 (catalog number AB9094-I, MilliporeSigma), mouse monoclonal IgG1 anti-ATP1A3 antibody, clone XVIF9- G10, dilution 1:2,000 (catalog number XVIF9-G10, Thermo Fisher Scientific), rabbit polyclonal IgG anti-TRPV2 antibody, dilution 1:250 (catalog number NBP1-32096, Novus Biologicals, Centennial, CO, USA), rabbit polyclonal IgG anti-PIKFYVE antibody, dilution 1:2,000 (catalog number 13361-1-AP, Proteintech, Rosemont, IL, USA), rabbit polyclonal IgG anti-CHOP antibody, dilution 1:1,000 (catalog number 15204-1-AP, Proteintech), mouse monoclonal IgG2b anti-beta Actin antibody, clone BA3R, dilution 1:10,000 (catalog number BA3R, Thermo Fisher Scientific).

#### Secondary antibodies

goat anti-mouse IgG (H+L)-HRP-conjugated antibody, dilution 1:5,000 (catalog number 1721011, Bio-Rad, Hercules, CA, USA), goat anti-rabbit IgG (H+L)-HRP-conjugated antibody, dilution 1:5,000 (catalog number 1721019, Bio-Rad), goat anti-rabbit IgG (H+L) cross-adsorbed secondary antibody with Alexa Fluor 647, dilution 1:400 (catalog number A-21246, Thermo Fisher Scientific).

### Plasmids

All site-directed mutagenesis (SDM) reactions were done with the Q5 Site-Directed Mutagenesis Kit (catalog number E0554S, New England Biolabs, Ipswich, MA, USA) according to the manufacturer’s protocol and sequence verified by Sanger sequencing. Plasmids were transformed into the NEB 5-alpha Competent *E. coli* (catalog number C2987H, New England Biolabs) and grown at 37°C unless specified otherwise. All plasmids were isolated with the PureLink HiPure Plasmid Maxiprep Kit (catalog number K210006, Thermo Fisher Scientific) before being used for transfections.

To direct the enhanced green fluorescent protein (EGFP) to the ER, the ER signal peptide (sequence: MKLGRAVLGLLLLAPSVVQA) was inserted by site-directed mutagenesis (forward primer: 5’- CTGGCGCCGTCCGTGGTGCA GGCGGTGGCTAGCGTGAGCAAGGGCGAGGAG- 3’, reverse primer: 5’-CAGCAGCAGGCCCAGCACGGCCCGGCCCAGCTTCATGGTGGCGGATCCGGT-3’) between the start codon and the EGFP coding sequence of an expression plasmid, which was itself derived from a tau1-441-EGFP plasmid, described previously ^35^. Subsequently, the four-amino-acid ER retention sequence (KDEL) was inserted immediately before the stop codon (forward primer: 5’-GAGCTGAAAGCGGCCGCTAGGCCT-3’, reverse primer: 5’- GTCCTTTTGTACAGCTCGTCC ATGCCG-3’), generating the spEGFP^KDEL^ plasmid.

The DHCR7 plasmid (catalog number RC200480) was obtained from Origene (Rockville, MD, USA). To create the spEGFP^KDEL^-P2A-DHCR7 plasmid, two restriction enzyme sites were introduced flanking the spEGFP^KDEL^ (EcoRI, catalog number R3101S, and SalI, catalog number R3138S, New England Biolabs) and the DHCR7 sequence (Avr2, catalog number R0174S, and MluI, catalog number R3198S, New England Biolabs) using SDM. Following this, a restriction digest was performed to isolate the fragments, which were then ligated using T4 DNA ligase (catalog number M0202S, New England Biolabs) into a doxycycline-inducible empty expression cassette, pCW57-MCS1-P2A-MCS2 (catalog number 89180, Addgene).

Plasmids coding for the open reading frame (ORF) of various proteins: ATP1A1 (catalog number OHu15960), ATP1A2 (catalog number OHu16492), ATP1A3 (catalog number OHu26791), ATP12A (catalog number OHu04875), ATP4A (catalog number OHu25737), NCX (catalog number OHu26285C), TRPV2 (catalog number OHu20085D) were obtained from Genscript (Piscataway, NJ, USA). Customized plasmids were also obtained from Genscript. These included the insertion of fluorescent proteins in ATP1A2-P2A-EGFP, NCX-P2A-mCherry, and TRPV2-c-EGFP. The ATP1A2-P2A-spEGFP^KDEL^ plasmid was generated by undertaking the same SDM reactions as for spEGFP^KDEL^ to insert the SP and KDEL sequences. Plasmids for mutant proteins ATP1A1^D811A^ (forward primer: 5’-CTCTGCATTGCCTTGGGCACTG-3’, reverse primer: 5’-GATGGTGACAGTCCCCAG-3’), ATP1A2^T378N^-P2A-spEGFP^KDEL^ (forward primer: AAGACGGGCAACCTCACCCAG-3’, reverse primer: 5’- GTCCGAGCAGATGGTGGAC-3’) and ATP1A2^L764P^-spEGFP^KDEL^ (forward primer: 5’- GAGGGCCGCCcGATCTTTGAC-3’, reverse primer: 5’- CTCCACCCCCGTGACGAT-3’) were created by SDM. TRPV2-NT (1-390) was created by a SDM deletion reaction (forward primer: 5’-ACTAGTCCAGTGTGGTGG-3’, reverse primer 5’-CTTGGGGATGAGCAGATCC-3’).

### siRNAs

All siRNAs were diluted in nuclease-free water to a stock concentration of 10 µM. ATP1A1 siRNA (catalog number s1720) was obtained from Thermo Fisher Scientific. ATP1A2 siRNA pools A and B (two different batches, catalog number sc-42660) as well as the three individual siRNA duplexes from pool A (catalog numbers sc-42660a, sc-42660b and sc-42660c) were purchased from Santa Cruz Biotechnology (Dallas, TX, USA). ATP1A2 siRNA pool C (catalog number M-006112-00- 0010) was purchased from Horizon Discovery (Cambridge, UK). ATP1A3 siRNA (catalog number sc-36012) and scrambled siRNA (catalog number sc-37007) were also obtained from Santa Cruz Biotechnology.

### rAAV vectors

#### CBh spEGFP^KDEL^ scAAV construct assembly

The CBh promoter and self-complimentary ITR AAV backbone plasmid was deposited to Addgene by the Michael J Fox Foundation MJFF (catalog number 194245, Addgene). The EGFP sequence was PCR-amplified from its donor plasmid (catalog number 22875, Addgene) in Q5 Hot Start High Fidelity 2X Master Mix (catalog number M0494, New England Biolabs) according to manufacturer instructions (forward primer: 5’- CAGGTTGGACCGGCTAGCACCGGTGCCACCATGAAGCTGGGCCG-3’, reverse primer: GTAATCCAGAGGTTGATTAGGAAGTTCATCCTTCTACTTGTACAGCTCGTCCATGCCG-3’). The backbone vector was digested with AgeI-HF and StuI (catalog numbers R3552 and R0187, New England Biolabs) simultaneously according to manufacturer instructions. This removed the ‘null’ sequence and linearized the plasmid. Then, the backbone was purified on a 0.9% agarose gel. The EGFP gene was inserted into the linearized plasmid using NEB HiFi Assembly 2X Master Mix (catalog number E2621, New England Biolabs) relying on overlap regions inserted at the ends of each PCR primer. To produce a rAAV vector coding for spEGFP^KDEL^, the SP and KDEL sequences were inserted into the pscAAV-CBh-EGFP plasmid using the same SDM reactions as mentioned above. This final plasmid was transformed into the NEB stable competent *E. coli* (catalog number C3040H, New England Biolabs) and grown at 30°C to minimize inadvertent recombination of its inverted terminal repeats (ITRs). The finalized plasmid was purified and sequence verified.

#### rAAV production and purification

The production and purification of rAAV vectors was based on a previously reported method [38]. Briefly, HEK293T cells were grown in a HYPERFlask with 1,720 cm2 surface area and 560 mL growth medium (CLS10031-4EA, MilliporeSigma). At 70- 80% confluency, the cells were triply transfected with the rAAV transfer plasmid, the rAAV packaging plasmid coding for Rep and Cap genes (catalog number 103005, Addgene), which we modified by SDM to code for the 9P31 capsid, and the helper plasmid coding for adenovirus genes E2A, E4 and VA (catalog number 112867, Addgene). Five days later, 3 mL Triton-X 100 (catalog number X100-100ML, MilliporeSigma), 250 µL RNAse A (catalog number EN0531, Thermo Fisher Scientific), 56 µL Pluronic F-68 (catalog number 24040-032, Thermo Fisher Scientific), and 56 µL turbonuclease (catalog number T4330, MilliporeSigma) were added to the cell culture medium in the HYPERFlask. This was followed by agitation of the flask at 150 rpm and 37°C for 1 hour. The cell lysates were then removed from the HYPERFlask, before rinsing the HYPERFlask with 140 mL of PBS. The lysate and PBS wash were then pooled and spun down at 4,000 g for 30 minutes to create a pellet of cell debris. The centrifugation supernatant was filtered through a 0.45 µm PES Autofil bottle top vacuum filter assembly of 1L (catalog number 1143-RLS, Foxx Life Sciences, Londonderry, NH, USA).

The filtrate was subsequently loaded onto a POROS GoPure AAVX pre-packed column (catalog number A36652, Thermo Fisher Scientific) at a rate of 0.2-0.3 mL/min over 44 hours using an ӒKTA fast protein liquid chromatography (FPLC) system (GE Healthcare, Chicago, IL, USA). Before eluting the rAAV, the affinity matrix was subjected to four washes at a rate of 0.5 mL/min as follows: 15 mL of 1× Tris-buffered saline (TBS) (0.05 M Tris/HCl, pH 7.6, 0.15 M NaCl), 15 mL of 2× TBS, 20 mL of 1x TBS with 20% ethanol (v/v), and 15 mL of 1× TBS. 12 mL of elution buffer consisting of 0.2 M glycine and 0.01% Pluronic F-68, pH 2-2.5, was used to elute the rAAV vectors across three 4 mL fractions, each collected in a tube containing 420 µL of neutralization buffer (1 M Tris/HCl, pH 8, 0.1% Pluronic F-68). Prior to use, each buffer was filtered through a 0.22 µM PES membrane with a Stericup Quick Release-GP filtration system (catalog number S2GPU11RE, MilliporeSigma).

The three elution fractions were then pooled and diluted with final formulation buffer (0.001% Pluronic F-68, 35 mM NaCl, and 1× PBS) to a total volume of 50 mL. The rAAV vectors were then filtered through a 0.22 µM PES Millex-GP Syringe Filter Unit (catalog number SLGPR33RS, MilliporeSigma) before concentration with an Amicon stirred cell ultrafiltration unit (catalog number UFSC05001, MilliporeSigma) with a 100 kDa Ultrafiltration Disc (catalog number PLHK04310, MilliporeSigma). The Amicon stirred cell ultrafiltration unit was attached to a nitrogen tank and placed on a magnetic stir plate set to a low to medium speed. The volume of the AAV vector concentrate was allowed to reach 5 mL before the Amicon stirred cell ultrafiltration unit was refilled with final formulation buffer to 50 mL. This step was repeated three times in total, and on the last concentration the contrate was reduced to 1 mL. The rAAV vectors were then removed from the Amicon stirred cell ultrafiltration unit with a pipette for storage in 20-50 µL aliquots at - 80°C. Any glass or plasticware in contact with rAAV vectors was coated with 0.005% Pluronic F- 68, 35 mM NaCl, 1× PBS before use.

#### rAAV titration

The titre of rAAV vector preparations was determined by qPCR using a published method [361]. Briefly, we prepared DNA standards by linearizing 20 µg of the respective transfer plasmid with the restriction enzyme ScaI (catalog number R3122S, New England Biolabs), then ran the product on a 0.9% agarose gel. A DNA cleanup kit (catalog number T1030S, New England Biolabs) was used to remove impurities from the digested DNA, before eight serial dilutions of this standard stock were completed in triplicate using UltraPure water.

When preparing rAAV vector preparations for titration, we initially digested DNA contaminants by DNAse I (catalog number EN0521, Thermo Fisher Scientific) at 37°C for 1 hour in digestion buffer (2 mM CaCl2, 10 mM Tris-HCl, and 10 mM MgCl2), using 5U of DNAse I added to 2 µL of the virus preparation in a 100 µL reaction. Next, DNAse I was deactivated by adding 5 µL EDTA (0.5 M, pH 8.0) (catalog number 15575-038, Invitrogen) to each tube before incubating for 10 minutes at 70°C. Next, the capsids of virus particles were digested with Proteinase K (catalog number PRK403.100, BioShop) to release viral genomes. This digestion was accomplished by adding 120 µL of buffer composed of 1 M NaCl and 1% (w/v) N-lauroylsarcosine sodium salt and 12 µg of Proteinase K to the DNAse I digestion product and incubating the reaction at 50°C for at least 2 hours. Proteinase K was then deactivated for 10 minutes at 95°C. The released viral genomes were then diluted 1:300 in UltraPure water and qPCR reactions were conducted in triplicate, each containing the 12.5 µL SYBR Green Master Mix (catalog number A46012, Thermo Fisher Scientific), 9.5 µL UltraPure water, 0.5 µL each of the forward and reverse primer at 2.5 µM initial concentration (forward primer: 5’-cctattgacgtcaatgacgg-3’, reverse primer: 5’-gatgtactgccaagtaggaaag-3’), and 2 µL of the diluted sample. A LightCycler 480 Instrument II (Roche Life Science Solutions, Basel, Switzerland) was used to denature DNA initially for 10 minutes at 95°C, then amplify the PCR target sequence during 40 cycles of 15 seconds at 95°C and 60 seconds at 60°C.

### Cell culture

#### Cell lines and maintenance

U2OS human osteosarcoma cells (catalog number HTB-96), HEK293 human embryonic kidney cells (catalog number CRL-1573), and HEK293T cells (catalog number CRL-3216) were acquired from American Type Culture Collection (ATCC) (Manassas, VA, USA) and cultured in Dulbecco’s Modified Eagle Medium (DMEM) with high glucose and sodium pyruvate (catalog number 11995065, Thermo Fisher Scientific) supplemented with 10% fetal bovine serum (FBS) (catalog number 12483020, Thermo Fisher Scientific), 2 mM GlutaMAX (catalog number 35050061, Thermo Fisher Scientific) and 50 U/mL Penicillin-Streptomycin (catalog number 15070063, Thermo Fisher Scientific). Cells were cultured in a humidified incubator at 37°C with 5% CO2 and were subcultured every 2-3 days at a 1:10 dilution.

#### Primary neuronal cultures

Primary neuronal cultures from ATP1A1^s/s^ mice were harvested from mouse embryos at P15-P17 of gestation. 6 cm Nunc EasYDish dishes (catalog number 150462, Thermo Fisher Scientific) were coated overnight before the harvest with 3 mL of 50 μg/mL poly-D-lysine hydrobromide (catalog number P6407, MilliporeSigma). The next morning, the coated plates were rinsed with sterile distilled water four times and air-dried in the biosafety cabinet. To collect neurons, a pregnant *ATP1A1^s/s^* mouse was sacrificed by cervical dislocation. Next, its embryos were carefully removed from the placenta and immediately placed into ice-cold DMEM to preserve tissue integrity. Next, the brains were extracted from the embryos, and the cerebellum and meninges were removed under a dissecting microscope (Model MS5-PS, Leica Microsystems, Wetzlar, Germany). The cortices were digested with a mix of 0.01% (w/v) papain (catalog number P5306, MilliporeSigma), 0.1% (w/v) neutral protease (catalog number P4630, MilliporeSigma), and 0.01% (w/v) DNase I (catalog number DN25, MilliporeSigma) for 25 minutes under gentle shaking at 37°C to ensure full dissociation of cells. Once the digestion was complete, the cortices were mechanically dissociated by gentle pipetting. Initially, a 5 mL pipette was used, followed by a P1000 pipette, and finally a glass-fired Pasteur pipette, to achieve a finer dissociation. After this process, the cell suspension was centrifuged at 300g for 5 minutes. Pelleted cells were resuspended in DMEM with 50% FBS and plated at a density of 5x 10^5^ per cm^2^. Two hours post-plating, the medium was replaced with fresh neurobasal medium (catalog number 21103049, Thermo Fisher Scientific), supplemented with 1x B27 (catalog number 17504044, Thermo Fisher Scientific), 2 mM GlutaMAX, and 100 U/mL penicillin and streptomycin. On the third day after plating, 1 μM cytosine β-D-arabinofuranoside (AraC) (catalog number C1768, MilliporeSigma) was added to the cultures to inhibit glial cell proliferation.

#### Transfections

For plasmid transfections, U2OS cells or HEK293 cells were detached with 0.25% trypsin-ethylenediaminetetraacetic acid (EDTA) (catalog number 25200056, Thermo Fisher Scientific) and seeded at 80% confluency in a 24-well glass bottom plate (catalog number P24- 1.5P, Cellvis, Mountain View, CA, USA) the day before transfection. For each well, 0.5 µg of plasmids were transfected with the Lipofectamine 3000 transfection reagent (catalog number L3000008, Thermo Fisher Scientific) according to the manufacturer’s protocol, with 1 µL of Lipofectamine 3000 and 1 µL of P3000 used. For siRNA transfections, cells were seeded at 50% confluency in a 24-well glass bottom plate and siRNAs were transfected at a final concentration of 30 nM with 1.5 µL of the Lipofectamine RNAiMAX transfection reagent (catalog number 13778075, Thermo Fisher Scientific) according to the manufacturer’s protocol. For western blot analysis, transfections were carried out in 6 cm plates.

### Cell treatments

#### Cardiac glycoside (CG) treatments

Primary cortical neuronal cultures from *ATP1A1^s/s^* mice were treated by replacing 50% of the cell culture media with pre-warmed media containing the specific concentrations of ouabain. Images were taken 6 hours after the treatment. For U2OS cells, strophantidin was added 6 hours after ATP1A2 plasmid transfection by replacing the entire transfection media with pre-warmed growth media containing the CG concentrations indicated in the text.

#### OphA and CsA treatments

One day before treatment with OphA or CsA, U2OS cells were detached with trypsin-EDTA and seeded at 80% confluency in either a 24-well glass bottom plate (for imaging experiments) or 6 cm plates (for western blot analyses). The plates were gently rocked in a crosswise manner to ensure even distribution of cells. The next day, OphA or CsA was added by replacing the existing media with pre-warmed media containing these compounds at concentrations indicated in the text. Tranilast, BAPTA-AM, and DIDS were mixed in the same tube as OphA at concentrations indicated in the text. Cells were imaged 3.5 hours after OphA treatment and 20 hours after CsA treatment.

#### Treatment in conjunction with plasmid transfections

For spEGFP^KDEL^ transfected cells, OphA, CsA, tranilast or a combination of OphA and tranilast were mixed and added the same way as described above 24 hours after the transfection. For TRPV2-c-EFGP and ATP1A2-P2A- spEGFP^KDEL^ transfected cells, tranilast was added 6 hours after the transfection the same way as described above.

#### Ca^2+^ indicators

U2OS cells were prepared the same way as previously mentioned for OphA treatment. 3 hours after OphA treatment, cells were incubated in 2 µM of Fluo-3 or 10 µM of Mag-Fluo-4 for 30 mins in Hanks’ Balanced Salt Solution (HBSS) (catalog number 14025092, Thermo Fisher Scientific). Cells were then washed two times with HBSS and kept in HBSS for de-esterification for an additional 15 mins before imaging.

### Animals

#### Animal source and husbandry

The study made use of *Atp1a1^s/s^* mice, which had been described before ^36^ and were kindly provided to us by the late Jerry B. Lingrel, University of Cincinnati College of Medicine. *C57BL/6J* mice (catalog number 000664) were purchased from The Jackson Laboratory (Bar Harbor, ME, USA). A maximum of five mice were housed in a single cage and cages were changed at least once every week. The mice were on a 12-hour light/dark cycle and were subjected to daily monitoring of health and activities. Mice were fed with 18% protein chow and provided with acidified water *ad libitum*. All animal procedures were in accordance with the Canadian Council on Animal Care, reviewed and authorized by the University Health Network Animal Care Committee and approved under Animal Use Protocol 6325.7.

#### Acute CG injections

Three-month-old wild-type (wt) or *Atp1a1^s/s^* mice were anesthetized by inhalation of 5% isoflurane and injected with 20 µM ouabain in phosphate-buffered saline (PBS) (catalog number 14190144, Thermo Fisher Scientific) via a free-hand intracerebral injection. 6 hours following the injection, mice were sacrificed and analyzed as indicated below.

#### Prion inoculation

Three 2-month-old *C57BL/6J* mice were inoculated with 30 µL of 1% (w/v) brain homogenate in PBS + 5% BSA (w/v) via intracerebral injections into the right parietal lobe. The source homogenate came from a *C57BL/6J* mouse that had succumbed to prion disease after being inoculated itself with Rocky Mountain Laboratory (RML) prions. Three negative control *C57BL/6J* mice were inoculated in the same manner with 30 µL of PBS + 5% BSA. Around 160 days post-inoculation (dpi), when the prion-inoculated mice reached the humane endpoints (two or more clinical signs of prion disease or >20% weight loss), the mice were sacrificed as described below.

#### AAV injections

90 days postinoculation (dpi), prion-inoculated and vehicle-inoculated control *C57BL/6J* mice were administered 1×10^12^ vg of scAAV-9P31-CBh-spEGFP^KDEL^ diluted in PBS to a total volume of 100 µL by single retro-orbital sinus injection on the left side using a 28 gauge insulin syringe (catalog number 329420, BD Canada, Mississauga, ON, Canada).

### Brain tissue analyses

#### Immunohistochemistry

Mouse brains were transcardiacally perfused with PBS, rapidly dissected and stored in 10 mL of neutral buffered formalin (catalog number HT501128-4L, Sigma) for 48 h, followed by immersion in 10 mL of 70% ethanol in water (v/v) for at least 7 days, before embedding in paraffin. 5 µM thick brain sections were cut from the paraffin block then mounted on positively charged glass slides. To deparaffinize and rehydrate the sections, they were immersed for 3 minutes each in three consecutive xylene baths, two 99% ethanol baths, one 95% ethanol bath, one 70% ethanol bath, followed by immersion in water. To retrieve the NeuN antigen, the slides were immersed in Tris-EDTA retrieval buffer, then subjected for 15 minutes to 115 °C in a pressure cooker and allowed to cool on the benchtop for 20 minutes, before being rinsed with distilled water for 5 minutes. Next, excess liquid was blotted off the slides with Kimwipes, sections were circled with a hydrophobic barrier pen and covered in blocking buffer for 1 h in a humidified chamber. Incubations with primary antibody occurred in blocking buffer overnight at 4 °C, followed by three 10-minute washes, 1 h incubation with secondary antibody (1:1,000 diluted) in blocking buffer, and two 20-minute washes. Next, mounting medium was applied in a straight line to one side of the sections, coverslips were placed such that they touched these lines and were gently lowered over the slide in a manner that forced the mounting medium evenly across the sections.

#### Cryo-sectioning of mouse brains for direct fluorescence detection

Mice were euthanized under deep anesthesia by inhalation of 5% isoflurane along with 1.5 L/min of oxygen. Brains were immediately removed from the brain cavity and meninges carefully removed. Brains were then transferred to a 15 mL tube containing 30% sucrose solution in PBS for cryoprotection and incubated at 4°C for up to 1.5 days, during which they sank to the bottom of the tube. Brains were then equilibrated in a 1:1 (v/v) mixture of 30% sucrose and Tissue-Tek O.C.T. Compound (catalog number 25608-930, VWR, Radnor, PA, USA) for 1 hour 4°C degrees. Next, brains were positioned in a cryomold (catalog number 70182, Electron Microscopy Sciences, Hatfield, PA, USA) and embedded in O.C.T for another 1 hour at 4°C degrees. Finally, the brains were frozen by partially immersing the cryomolds in a liquid nitrogen-chilled 2-methylbutane bath for 2-3 mins until the O.C.T completely froze. The frozen blocks were placed on dry ice to briefly air-dry the residual 2-methylbutane before being transferred to a −80°C freezer for storage.

Prior to cryo-sectioning, the brains were allowed to warm up in the Cryostat (model HM525 NX, Thermo Fisher Scientific) set at −21°C for at least 2 hours. 40 µM sections were cut and collected on a piece of adhesive Cryofilm (catalog number C-FUF303, Section-Lab Co. Ltd., Yokohama, Kanagawa Prefecture, Japan) using Kawamoto’s Film Method ^37^. Sections were then washed for 5 mins in PBS and mounted with mounting media containing DAPI (catalog number ab104139, Abcam) before imaging.

### Western blotting

U2OS cells were rinsed with ice-cold PBS and lysed in ice-cold lysis buffer for 5 mins on ice. The lysis buffer consisted of 1% (w/v) NP-40, 150 mM NaCl, 150 mM Tris/HCL, pH 8.3, as well as a protease inhibitor cocktail (catalog number 11836170001, MilliporeSigma). Cell lysates were then harvested with a cell scraper and insoluble cell debris pelleted by 5-minute centrifugation at 21,600 g at 4 °C. The cleared supernatant was transferred to a separate tube and the protein concentration measured by the bicinchoninic acid assay (BCA) using the Pierce BCA Protein Assay Kit (catalog number 23225, Thermo Fisher Scientific). Subsequently, protein concentrations were adjusted and 45 µL of sample (containing 60 µg of protein) were diluted with 15 µL of 4X LDS sample buffer (catalog number 84788, Thermo Fisher Scientific) containing 2.5% β-mercaptoethanol (v/v) to achieve a final protein concentration of 1 µg/uL. The sample was then heated at 70°C for 10 mins. If not used analyzed by sodium dodecyl-sulfate polyacrylamide gel electrophoresis immediately, samples were snap-frozen in liquid nitrogen and stored in −80°C.

SDS-PAGE analyses relied on 4–12% Bolt Bis-Tris gels (catalog number NW04120BOX, Thermo Fisher Scientific) using 1x MOPS SDS running buffer (catalog number J62847.AP, Thermo Fisher Scientific). The gels were immersed in a Mini Gel Tank (catalog number A25977, Thermo Fisher Scientific) and proteins separated at 165 volts for 40 minutes. Protein bands were transferred to 0.45 μm polyvinylidene fluoride membranes (catalog number IPVH00010, MilliporeSigma) using a Tris-Glycine transfer buffer that contained 25 mM Tris, 192 mM glycine, and 20% (v/v) methanol. The transfer was done during 1 hour at 35 V. Next, the PVDF membrane underwent blocking in 10% skimmed milk diluted in Tris-buffered saline (25mM Tris, 150mM NaCl) with 0.1% Tween-20 (TBST) (catalog number TWN508, BioShop) for 1 hour at room temperature. Subsequently, the membranes were incubated with gentle rocking in primary antibody solutions diluted at the indicated dilution factors in 5% skimmed milk at 4°C overnight. The following day, the membranes underwent three 5-minute washes in TBST with moderate shaking, followed by 1-hour incubation in the respective secondary antibody solutions in 5% skimmed milk at room temperature. Following another three 5-minute washes in TBST, the membranes were incubated for 90 seconds in 1 mL of enhanced chemiluminescence (ECL Pro) reagent (catalog number NEL122001EA, PerkinElmer Health Sciences Canada Inc., Woodbridge, ON, Canada). To visualize the signals, the membranes were exposed to autoradiography film (catalog number MED-CLMS810, Mandel Scientific, Toronto, ON, Canada) and developed with a film developer (Model SRX-101A, Konica-Minolta).

### Immunocytochemistry

Cells were washed 1–2 times with PBS and subsequently fixed with 4% formaldehyde (FA) diluted from a 36.5-38% stock (catalog number F8775-500ML, MilliporeSigma) for 10–20 minutes at room temperature. After fixation, cells were washed three times with PBS and stored in PBS at 4°C until further use. For permeabilization, cells were incubated in a buffer containing 1% (w/v) bovine serum albumin (BSA) (catalog number ALB001.50, Bioshop Canada, Burlington, ON, Canada), 0.1% Triton X-100, and 0.3M glycine (catalog number GLN001.1, Bioshop Canada) in PBS for 20 minutes. Primary antibodies were then incubated in PBS with 1% BSA overnight at 4°C, using the indicated dilutions. The next day, cells were washed three times with PBS and then incubated with the Alexa Fluor secondary antibody at the indicated dilutions in PBS with 1% BSA for 90 minutes at room temperature, while protected from light. After three additional PBS washes, samples were mounted using ProLong Gold Antifade Mountant with DAPI (catalog number P36935, Thermo Fisher Scientific) and imaged.

### Microscopy

Cell grown in culture were imaged on 24-well glass bottom plates with the Zeiss AXIO Observer 7 Inverted LED Fluorescence Microscope (Carl Zeiss Canada Ltd., Toronto, ON, Canada). Cryosections of brain samples were imaged on Fisherbrand Superfrost Plus Microscope Slides (catalog number 22-037-246, Thermo Fisher Scientific) under #1.5 glass cover slips (catalog number 48393-060, VWR) with the Zeiss LSM 880 inverted confocal microscope (Carl Zeiss Canada Ltd.).

### Statistical analyses

Comparisons of percentages of cells exhibiting PNS dilations were undertaken by tallying vacuolated versus total cells in fields of views in three biological replicates. Densitometry analyses of western blot signals of three biological replicates were done with Image J. P-values were calculated in Excel using the two-sample t-test assuming equal variance. The significance threshold is set at p < 0.05. One (*), two (**), and three (***) asterisks signify p-values of <0.05, <0.01, and <0.001, respectively. When the p-value is higher than the threshold, the abbreviation ‘N.S’ is shown.

## RESULTS

### Acute toxic exposure of rodent brains to CGs causes spongiform neuronal degeneration in *Atp1a1^s/s^* but not in wild-type mice

We set out to clarify if the cell-type specific pattern of α1 subunit expression may underlie the selective brain vacuolation that had been reported in rodent astrocytes upon acute and toxic CG exposure. To this end, we bolus injected ouabain—the CG that was used in prior studies, intracerebrally into wild-type mice or *Atp1a1^s/s^* to mice that had been rendered sensitive to CGs by the introduction of two human amino acids known to control CG binding (**Fig 1A**). Six hours after these injections, mouse brains were formaldehyde perfused and post-fixed, then paraffin-embedded and immuno-stained with antibodies against the neuron-specific marker NeuN or counterstained with hematoxylin & eosin. Consistent with prior reports, these analyses revealed the normal morphology of cells within the hippocampal formation if brains were injected with vehicle phosphate buffered saline (**Fig 1B**) and a selective CG vulnerability of astrocytes in wild-type mice that spared neurons expressing the refractory endogenous α1 subunit (**Fig 1C**). In contrast, hippocampal sections from *Atp1a1^s/s^* mice exhibited both astrocytic and neuronal cellular pathology, characterized by condensed nuclei and abundant vacuolation (**Fig 1D**). The subsequent analysis of primary neural cultures of cells harvested from *Atp1a1^s/s^*mice corroborated the conclusion that toxic levels of CGs can cause vacuolation (**Fig 1E**), with prominent perinuclear vacuoles forming at 1 µM ouabain concentrations.

**Fig. 1.**
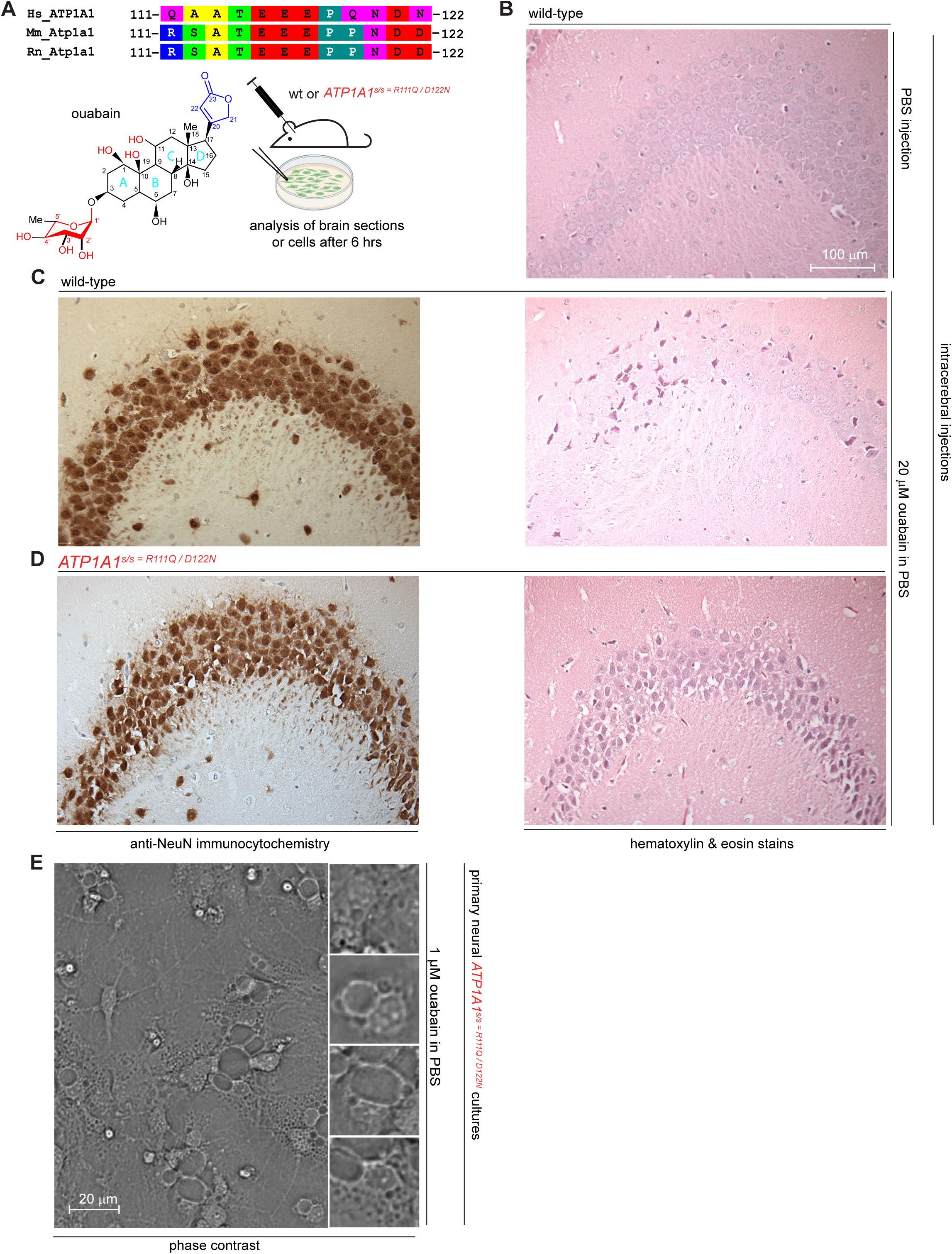
Acute toxic exposure of rodent brains to CGs causes spongiform neuronal degeneration in Atp1a1^s/s^ but not in wild-type mice. (A) Multiple sequence alignment of a short stretch of amino acids that contribute to the CG binding pocket in human, mouse, and rat NKA α1 subunits. The CG resistance of rodents has been ascribed to arginine and aspartic acid residues found in positions 111 and 122 of their α1 subunits^31^. The cartoon shown below the sequence alignment depicts the design of the acute CG exposure experiments undertaken with mice and cells in culture. (B) Hippocampal CA2/CA3 region of wild-type mouse that received a negative control PBS injection. (C) CG-poisoned hippocampal CA2/CA3 region of wild-type mice probed with an anti-NeuN antibody that labels neurons next to H & E stain. (D) CG-poisoned hippocampal CA2/CA3 region of *Atp1a1^s/s^* mice, analyzed as in Panel C. (E) Phase contrast images of primary neural cultures of *Atp1a1^s/s^*mice that were poisoned with 1 µM ouabain levels in their cell culture medium.

### Overexpression of NKA α subunits—not their suppression—causes profound dilation of the ER and PNS

To determine if CG exposure had caused vacuolation by inhibiting one or more of the NKA isoforms or through some other unknown mechanism, we obtained commercial small interfering RNAs (siRNAs) for the three human NKA α subunits. Looking for a human cell model that can be easily transfected and has been used for vacuolation studies before, we decided to transfect these silencing reagents into U2OS cells.

This experimental approach led us to chase initially a false lead. It will be briefly told here because lessons learned may be a valuable caution to others working with siRNA pools: When we transfected the siRNA pools into U2OS cells, we observed that the pool of siRNAs designed to target NKA α2 (*ATP1A2*) caused indeed a robust vacuolation phenotype (**S1 Fig**). Because the phenotype was characterized by a large vacuole adjacent to the nucleus, we suspected it might represent a dilation of the PNS. To investigate, we assembled a plasmid coding for an enhanced green fluorescent protein (EGFP) fused to an N-terminal ER signal sequence—borrowed from the ER-resident protein calreticulin—and a C-terminal ER retention motif ‘KDEL’ (spEGFP^KDEL^). Next, we co-transfected the siRNA pool targeting *ATP1A2* alongside this reporter plasmid and observed our hypothesis of the vacuoles representing dilations of the ER and PNS validated (**S1 Fig A**). Surprisingly, when we subsequently analyzed *ATP1A2* levels by western blotting, we did not find them to be reduced in samples that were derived from cells treated with the siRNA (pool A) targeting ATP1A2 (**S1 Fig B**). Considering that perhaps a non-RNA contaminant in pool A, rather than the siRNAs themselves, may be responsible for inducing the phenotype, we digested its siRNAs with RNAse A (**S1 Fig C**), then repeated the experiment and observed that the reagent no longer caused the dilation phenotype. Assured that the effect was RNA-dependent, we targeted *ATP1A2* with individual constituent siRNAs of pool A. We also purchased two separate siRNA pools B and C targeting *ATP1A2* transcripts from the same and a different vendor. Only pool A induced the ER/PNS dilation phenotype (not shown). Next, we analyzed RNA concentration-adjusted levels of the various *ATP1A2* siRNAs by agarose gel electrophoresis and observed visible levels of siRNAs in pool A to be lower than expected (**S1 Fig D**). Finally, we undertook RT-qPCR analyses and observed the original siRNA pool A had not lowered but increased ATP1A2 mRNA levels (**S1 Fig E**). We could not find any authoritative published records showing that partially degraded siRNAs stabilize the transcripts they are designed to degrade. Nevertheless, this became the most plausible interpretation of our experimental results.

Based on these data, we revised our hypothesis to propose that acute ouabain poisoning may trigger cells to increase their production of NKAs to protect themselves, leading to dilations of the ER and PNS. Consequently, we obtained overexpression constructs for the three NKA α subunits expressed in the brain and transfected them into the U2OS cell model. Validating the revised hypothesis, we observed vacuoles forming upon overexpression of all three NKA α subunits (**Fig 2A**). To determine whether the dilation phenotype correlates proportionally with the expression of the respective NKA α subunits, rather than being induced *in trans*, we next transfected U2OS cells with an expression plasmid that encoded both ATP1A2 and EGFP, linked by a self-cleaving P2A sequence. Vacuolation of U2OS cells transfected with this construct was observed to be proportional to the levels of the EGFP marker, indicating that the phenotype is generated in *cis* by the overexpression of the NKA α subunit (**Fig 2B**). The concomitant transfection of the spEGFP^KDEL^ reporter again confirmed the induced vacuoles to represent ER/PNS dilations (**Fig 2C**). Finally, to confirm the identity of the dilated subcellular compartment using an orthogonal approach, we used an antibody against the KDEL motif and found that the vacuoles induced by ATP1A2 overexpression were surrounded by the antibody’s immunocytochemical signal (**Fig 2D**). A body of literature has established a close functional relationship of NKAs and sodium calcium exchangers (NCX). We therefore investigated if an alteration of NCX levels might be able to induce the same phenotype, possibly reflecting this functional relationship. This was indeed the case, i.e., when we overexpressed NCX in U2OS cells, we also observed a significant increase in the percentage of U2OS cells exhibiting the PNS dilation phenotype (**Fig 2E, F**).

**Fig. 2.**
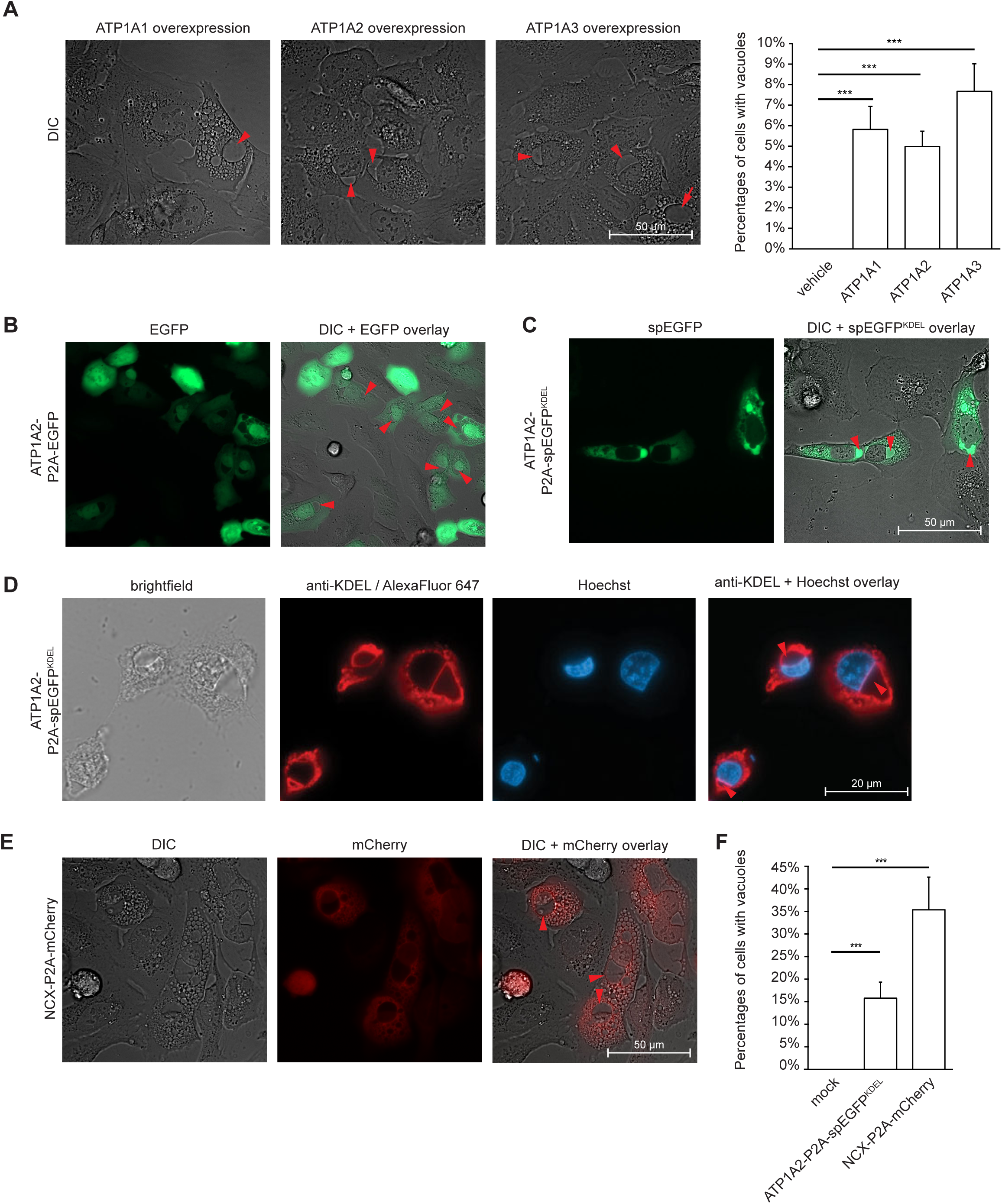
Overexpression of NKA α subunits causes profound ER/PNS dilations. (A) Transient overexpression of any of the three brain NKA α subunits in U2OS cells caused large cytosolic vacuoles that frequently abutted the nucleus. The percentages of cells which exhibited this phenotype are shown in the graph on the right. (B) Expression of ATP1A2-P2A-EGFP-KDEL construct documents that ER/PNS dilations manifest in transfected cells but not in non-transfected neighboring cells in trans. (C) Transfection of U2OS cells with plasmid coding for the overexpression of ATP1A2-P2A-spEGFP-KDEL established the identity of the vacuoles to be ER/PNS-derived by their green fluorescence. (D) Immunocytochemical analyses of U2OS cells with anti-KDEL antibody validates large vacuole adjacent to nucleus to represent dilated PNS. (E) Overexpression of NCX-P2A-mCherry in U2OS cells also gives rise to vacuolation. (F) Graph depicting percentage of cells exhibiting vacuoles after transfection with plasmids expressing ATP1A2-P2A-EGFP or NCX-P2A-mCherry. Where red or green arrowheads are used within the fluorescence or immunohistochemical analyses shown in this figure, they point toward PNS dilations.

### Overexpression of distant P-type ATPase paralogs is sufficient and ion transport function is dispensable for inducing ER/PNS dilation phenotype

Next, we investigated if the ER/PNS dilation phenotype can also be invoked by the overexpression of more distant paralogs of ATPases within the P2c branch of this gene family. ATP4A and ATP12A are the closest phylogenetic neighbors to NKA α subunits (**Fig 3A**). In contrast to NKAs, they pump H^+^ and K^+^ ions under ATP consumption and are also known as the gastric and non-gastric H^+^/K^+^-ATPases, respectively. Whereas ATP4A forms a heterodimeric complex with its designated β subunit (coded by *ATP4B*), ATP12A has been reported to pair with the NKA β1 subunit (coded by *ATP1B1*) ^38^. Overexpression of either protein in U2OS cells led to the same ER dilation phenotype (**Fig 3B**).

**Fig. 3.**
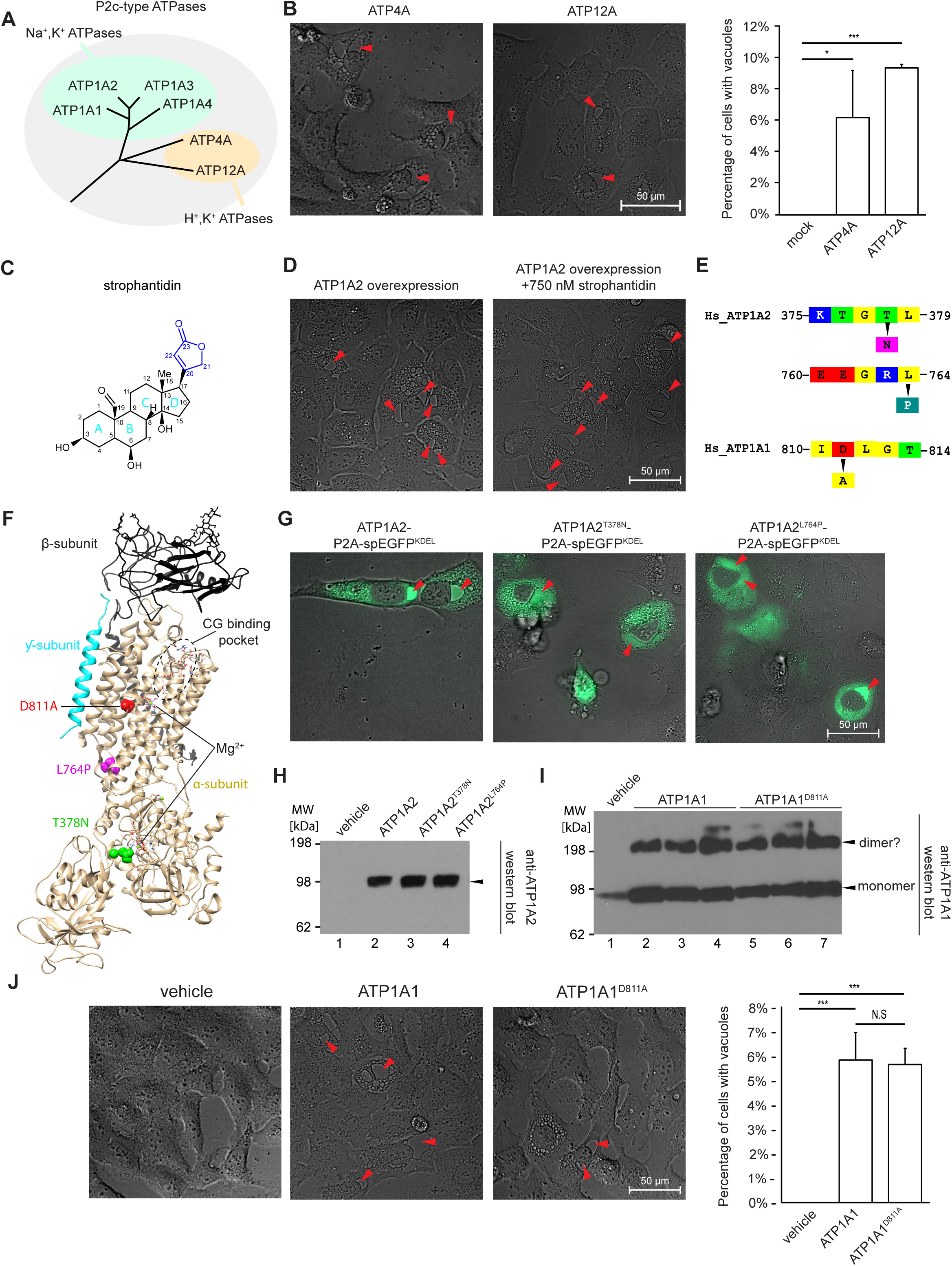
Overexpression of distant P-type ATPase paralogs is sufficient and ion transport function is dispensable for inducing ER/PNS dilation phenotype biology. (A) Dendrogram showing phylogenetic relationships amongst P2c-type ATPases. (B) Overexpression of ATP4A or ATP12A in U2OS also causes vacuoles. Graph depicting percentages of cells with vacuoles is shown on the right. (C) Chemical structure of strophantidin, a CG known to exhibit relatively high cell permeability due to it carrying a low number of hydrogen donors and being devoid of a carbohydrate moiety. (D) High concentrations of strophantidin cannot block the ATP1A2 overexpression-dependent vacuolation of U2OS cells. ATP1A2 overexpressing cells, side-by-side exposed to vehicle or 750 nM strophantidin, exhibit the characteristic ER and PNS dilations. (E) Subset of mutations in human NKA α subunits reported to ablate their ion channel functions, i.e., namely ATP1A2 mutations T376N and L764P and the ATP1A1 mutation D811A. (F) Location of selected mutations in NKA α subunits. Depicted is a model of human ATP1A1 rendered with UCSF Chimera (v 1.1.2, San Francisco, CA, USA) based on its homology to the 3.4 Å crystal structure of its porcine ortholog (PDB entry 4HYT), which exhibits 98.6% sequence identity. Note that the amino acid sequences of proteins expressed by ATP1A1 and ATP1A2 are 86.8% identical, enabling a close approximation of the location of the three mutations in this representative NKA α subunit. (G) Ion transport function is dispensable for ER/PNS dilation caused by ATP1A2 overexpression, because overexpression of functionally ablated mutants ATP1A2T378N or ATP1A2L764P also induces this phenotype in U2OS cells. (H) Overexpression of ATP1A2 and its mutant derivatives led to similar levels of expression of these heterologous proteins in U2OS cells. (I) Western blot depicting that the overexpression of ATP1A1 or its mutant ATP1A1^D811A^ strongly increased total steady state levels ATP1A1 in U2OS cells. (J) The functionally ablated mutant ATP1A1^D811A^ is equally potent in its ER/PNS dilation inducing properties as wild-type ATP1A1. A graph depicting the percentages of cells exhibiting vacuoles is shown on the right. Red arrowheads point toward PNS dilations.

Is the ion pump function of overexpressed P2c-type ATPases responsible for the formation of the ER/PNS vacuoles, for instance by promoting an ion imbalance across the lipid bilayer surrounding this compartment? To begin to address this question, we combined overexpression of the NKA α2 subunit with exposure of cells to strophantidin. Strophantidin was chosen because it lacks carbohydrates and carries fewer hydrogen donors than ouabain (**Fig 1A**), thereby facilitating its entry into cells to inhibit the ER-based NKAs (**Fig 3C**). These experiments did not show a convincing reduction in the levels of vacuolation (**Fig 3D**). We understood them to be flawed because acute CG exposure had led us to the ER/PNS dilation phenotype (**Fig 1**), and we suspected that strophantidin might not only inhibit the overexpressed α2 subunit itself but may also have inhibited other endogenous NKA α subunits with poorly understood consequences for their expression.

To address these caveats, we overexpressed NKA α subunits carrying mutations known to ablate the ion transport mechanism of NKAs in humans, namely ATP1A2^T378N^, ATP1A2^L764P^, and ATP1A1^D811A^ (**Fig 3E**). The choice fell on these mutations because they affect different aspects of the ion transport mechanism by targeting spatially distant sites in NKA complexes that comprise ATP1A2 or ATP1A1 subunits (**Fig 3F**). Whereas ATP1A2^T378N^, ATP1A2^L764P^ target the ATP binding site and a flexible hinge region within the NKA α subunit, the ATP1A1^D811A^ mutant perturbs Na^+^ ion coordination and has been reported to underlie a Charcot-Marie-Tooth Type 2 phenotype in humans carrying it ^39, 40^. When we transfected plasmids into U2OS cells that coded for wild-type ATP1A2 or the mutated NKA α subunits ATP1A2^T378N^ or ATP1A2^L764P^, we observed that these mutants retained the ability to induce the ER and PNS dilation phenotype (**Fig 3G**), despite steady-state protein levels not being affected by the mutations (**Fig 3H**). Similarly, when we introduced the ATP1A1^D811A^ levels, we observed that its expression was not diminished relative to wild-type steady-state ATP1A1 levels (**Fig 3I**). As for the ATP1A2 mutants, the microscopical analyses of cells transfected with the ATP1A1^D811A^ mutant revealed the ER dilation phenotype to be independent of the activity of this NKA α subunit (**Fig 3J**).

These results established that the overexpression of several paralogs of a family of ion transporters is sufficient to induce the ER/PNS dilation phenotype, and the ion transport capability of these transporters is not required for its manifestation.

### Establishment of alternative paradigms for studying the molecular mechanisms of ER and PNS dilation

A deep literature survey revealed that ER and PNS dilations, like the phenotype induced by overexpression of P2c-type ATPases, have been reported across various human cell models in response to range of cellular insults. This survey also uncovered inconsistency in the terminology used to describe these dilation phenotypes, with terms ranging from autosis, oncosis, and paraptosis, to necroptosis. After carefully examining this scattered body of work, we decided to adopt two additional ER and PNS dilation paradigms, as they provided more straightforward and simplified assays.

One was based on overexpression of 7-dehydrocholesterol reductase (DHCR7) ^41, 42^, which we selected for further studies because—in contrast to P-type ATPases—DHCR7 is not known to require other closely associated molecules for its function. To decouple cellular transfection from vacuolation and enable fluorescent visualization of vacuoles, we inserted DHCR7 into a doxycycline inducible cassette downstream of an spEGFP^KDEL^ coding sequence that it was linked to through a self-cleaving motif borrowed from porcine teschovirus-1 (P2A) (**Fig 4A**). Transfection of this expression plasmid into U2OS cells (**Fig 4B**), followed by doxycycline addition, caused ER and PNS dilations to form within 36 h (**Fig 4C**).

**Fig. 4.**
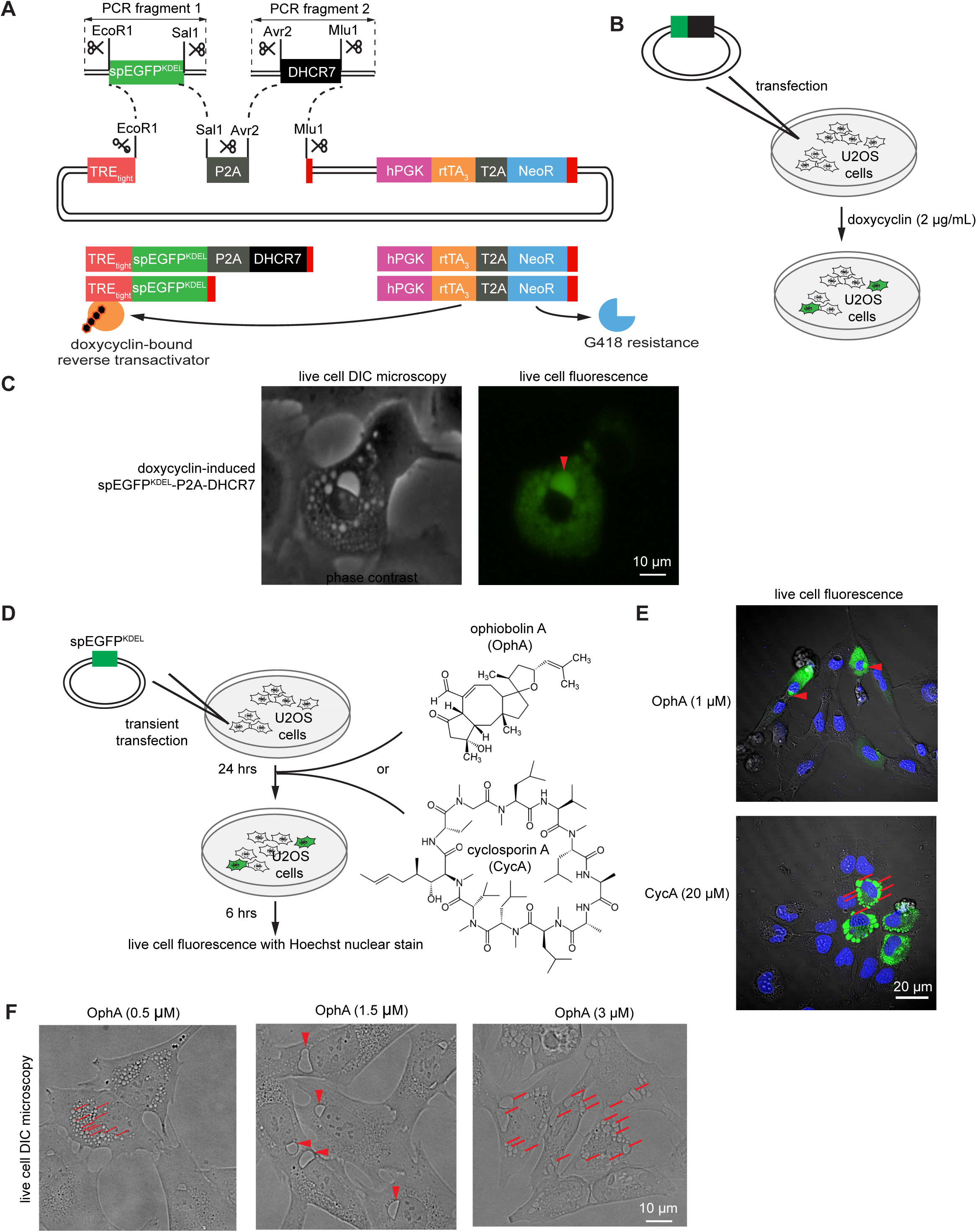
Establishment of alternative ER/PNS dilation paradigms for studying the underlying cellular biology. (A) Cartoon showing design of inducible expression constructs and the doxycycline-dependent transactivation of spEGFP or the concatemeric spEGFP-P2A-DHCR7 expression cassette. (B) Workflow of transfection and doxycycline-based induction of spEGFP in human U2OS cells. (C) Expression of DHCR7 causes the appearance of ER/PNS dilations in a subset of cells that can be identified by their spEGFP fluorescence. (D) Implementation of vacuolation paradigms induced by addition of ophiobolin A (OphA) or cyclosporin A (CsA), shown in their chemical structures to the right. (E) Evidence that 1 µM OphA or 20 µM CsA can induce ER/PNS dilations in U2OS cells. (F) Concentration-dependent presentation of vacuoles formed by OphA poisoning of U2OS cells. Note that at lower OphA concentrations (0.5 µM) many small vacuoles can be seen, whereas intermediate concentrations of OphA (1.5 µM) predominantly induce large PNS dilations that abut cellular nuclei, and cells exposed to high OphA concentrations (3 µM) are characterized by many large vacuoles. Red arrowheads, thick red lines, and thin red lines point toward representative PNS dilations, large cytoplasmic ER vacuoles, and small cytoplasmic ER vacuoles, respectively.

The second paradigm was based on a fungal phytotoxic metabolite, known as ophiobolin A (OphA) (**Fig 4D**) ^43^, whose addition to a variety of cell models had been described to induce ER and PNS dilations ^44, 45^. We also validated the potency of cyclosporin A to induce this phenotype ^46^ but did not pursue this line of investigation further, as we preferred the 20-fold higher potency of OphA for inducing the ER/PNS dilation phenotype. To facilitate assignment of the identity of vacuoles, the acute toxic addition of OphA was preceded by transfection with the spEGFP^KDEL^ reporter 24 hours earlier (**Fig 4E**). As anticipated, exposure of U2OS cells to OphA induced vacuolation within about 2 hours, with maximum dilation of the PNS reached after 4 to 6 hours (**Fig 4F** and **S2 Video**). In contrast to the protein overexpression paradigms, this pharmacological approach facilitated dose-response analyses. The latter revealed low concentrations of OphA to result in many small ER dilations, intermediate OphA exposure to cause the characteristic vacuoles we had observed in the protein overexpression paradigms, whose most striking feature were profound dilations of the PNS, and high levels of OphA to cause several large ER vacuoles (**Fig 4G**).

### Tranilast, an inhibitor of TRPV2 channels, blocks the OphA-induced PNS dilation

It has been proposed that increased protein density within the ER can give rise to an osmotic imbalance, which in turn may engender a net influx of water ^47^. Although conceptually straightforward, this notion is difficult to reconcile with the rapidity and absence of protein densities in the PNS swelling we observed. More plausible seemed that an ion imbalance generated the osmotic pressure, leading to rapid dilation and largely empty vacuoles. Having ruled out that the ion transport function of the NKAs itself is responsible (**Fig 3F**), we considered other known channels or pumps that control the ion balance of the ER and PNS. A breakthrough in this research occurred when we recognized the multi-modal characteristics of the transient receptor potential vanilloid channels (TRPV), and of features ascribed to one member of this family, TRPV2: 1) This channel was originally identified based on its responsiveness to insulin-like grow factor-1 (IGF1) stimulation ^48^, and we were aware that overexpression of the IGF1 receptor (IGF1R) had been shown to cause ER and PNS dilation ^49^. 2) TRPV2 channels interact with certain toxins, including cannabidiol ^50, 51^, which has been shown to induce ER swelling ^52^. 3) We also had noticed a paper reporting that overexpression of the TRPV2 paralog TRPV1, known as vanilloid receptor subtype 1, can induce a phenotype that the authors characterized as paraptosis—regrettably, no images were included in this prior report to assess if it has similarity to our phenotype ^53^. 4) A subset of TRPV channels have been reported to act as ER-based mechano-sensors, which we presumed make them good candidates for being involved in a phenotype characterized by osmotic swelling. 5) TRPV2 can also be activated by certain lipids, whose balance might be altered when members of the cholesterol synthesis pathway are overexpressed, as is likely the case in the DHCR7 overexpression paradigm ^41, 42^.

To begin to investigate the possible involvement of TRPV2 in the ER and PNS dilation phenotype, we treated cells with OphA in the presence or absence of 100 µM concentrations of tranilast, a known inhibitor of TRPV2 channels. The result was striking, as tranilast alone had no noticeable effect on the morphology of the cells but when added alongside OphA, it completely blocked the ER and PNS dilation phenotype (**Fig 5A**). Even 10 µM concentrations of tranilast in the U2OS cell culture medium were sufficient to reduce the size of the OphA-dependent swelling of the ER and PNS. This borderline effective tranilast concentration is mentioned here because it matches a previously reported ID50 for the tranilast-based inhibition of TRPV2 ^54^. This higher potency for TRPV2 sets this specific member of the vanilloid subfamily apart from other TRPV channels, which are known to be less sensitive to this agent (**Fig 5B**).

**Fig. 5.**
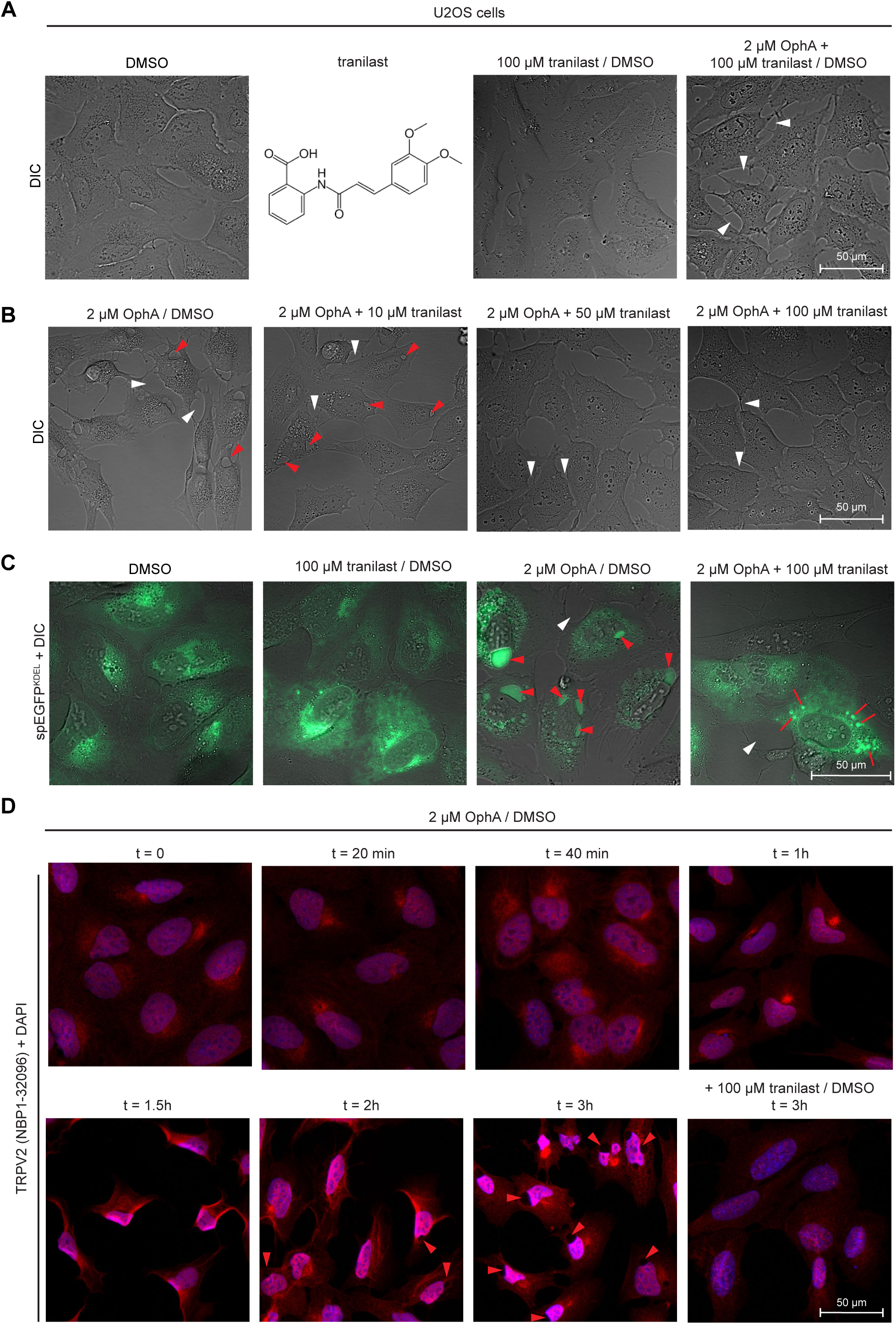
Tranilast, an inhibitor of TRPV2 channels, blocks the OphA-induced PNS dilation. (A) Exposure of U2OS cells to 100 µM tranilast does not cause any overt changes to their morphology but when combined with OphA addition to the cell culture medium, it blocks vacuolation. (B) Tranilast blocks the OphA-dependent vacuolation in a concentration dependent manner and mostly eliminates vacuoles from forming at 50 µM concentration. (C) In cells transfected with a plasmid coding for spEGFP, the tranilast-dependent inhibition of vacuolation leads to the formation of dense puncta that label with EGFP but are not recognizable as vacuoles by DIC microscopy. (D) Time course of TRPV2 signal distribution in cells exposed to OphA. One hour upon OphA addition to the cell culture medium of U2OS cells, TRPV2-immunocytochemical signals were observed to condense in a location adjacent to the nucleus after, then become more intense and spread to the plasma membrane during the next hour, followed by ER/PNS dilation at 3 hours. The addition of tranilast blocked the relocation of TRPV2 and the ER/PNS dilation. Red and white arrowheads point toward representative PNS dilations and intercellular spaces, respectively. Red lines indicate small ER vacuoles.

TRPV2 channels are understood to act as Ca^2+^ leak channels that localizes to the ER in unstimulated cells, raising the prospect that an increase in ER Ca^2+^ levels might occur when TRPV2 channels cannot open. Considering this context, we found it difficult to reconcile that U2OS cells exposed to tranilast alone did not develop ER/PNS dilations (**Fig 5A**). When we preceded the OphA and tranilast treatments with transfection of the spEGFP^KDEL^ reporter, we observed that instead of the large PNS dilations, the ER appeared to be condensed into compact, highly fluorescent dots (**Fig 5C**).

These observations raised the question of how tranilast inhibits TRPV2 function. Although the compound is marketed as a TRPV2 inhibitor, no specific binding pocket has been identified. Prior work suggested that other modes of action should be considered, including that it may impact the subcellular distribution of the channel ^55, 56^. To begin to answer this question, we treated U2OS cells with OphA and followed the subcellular localization of endogenous TRPV2 by immunocytochemistry. This experiment revealed the translocation of TRPV2 from the ER to the plasma membrane concomitant with an increase in steady-state TRPV2 levels approximately 1.5 hours after OphA addition to the cell culture medium, i.e., preceding the ER/PNS dilations by about 30 minutes. No such changes were seen in cells exposed to tranilast alone (**Fig 5D**).

These results support a model in which tranilast does not directly inhibit TRPV2, but instead prevents its upregulation and translocation to the plasma membrane, allowing the channel to maintain its function as an ER-based Ca^2+^ channel. Additionally, the results suggest that exposure to OphA leads to the temporary relocation of TRPV2 channels from the ER to the plasma membrane, which could enhance Ca^2+^ influx. This relocation may also increase osmotic Ca^2+^ pressure in the ER, owing to the disruption of the TRPV2 Ca^2+^ leak function.

### ER and PNS dilation phenotypes caused by *TRPV2* or *ATP1A2* overexpression can be blocked by tranilast

Since tranilast may have other cellular targets than TRPV2, we investigated if the mere overexpression of this channel protein can cause OphA-like swellings of the ER and PNS. We anticipated the overexpressed protein to increase Ca^2+^ entry into the cell because we assumed that the heterologous promoter may drive up TRPV2 levels artificially, thereby leading to overall increases in cellular Ca^2+^ levels that the additional ER-embedded TRPV2 Ca^2+^ leak channels may not be able to compensate. The situation may be exacerbated if the burden of the TRPV2 overexpression on the protein folding machinery in the ER would itself trigger the ER and PNS dilation phenotype we had seen when overexpressing other transmembrane proteins in U2OS cells. The results validated this hypothesis, showing profound ER/PNS dilation when we overexpressed a TRPV2 protein that was C-terminally fused to EGFP (TRPV2^EGFP^). When cells were exposed to both TRPV2^EGFP^ overexpression and OphA treatment, the phenotype became more pronounced, indicating a potential synergistic effect between these factors. However, this enhanced phenotype was still reversible with the addition of tranilast (**Fig 6A**).

**Fig. 6.**
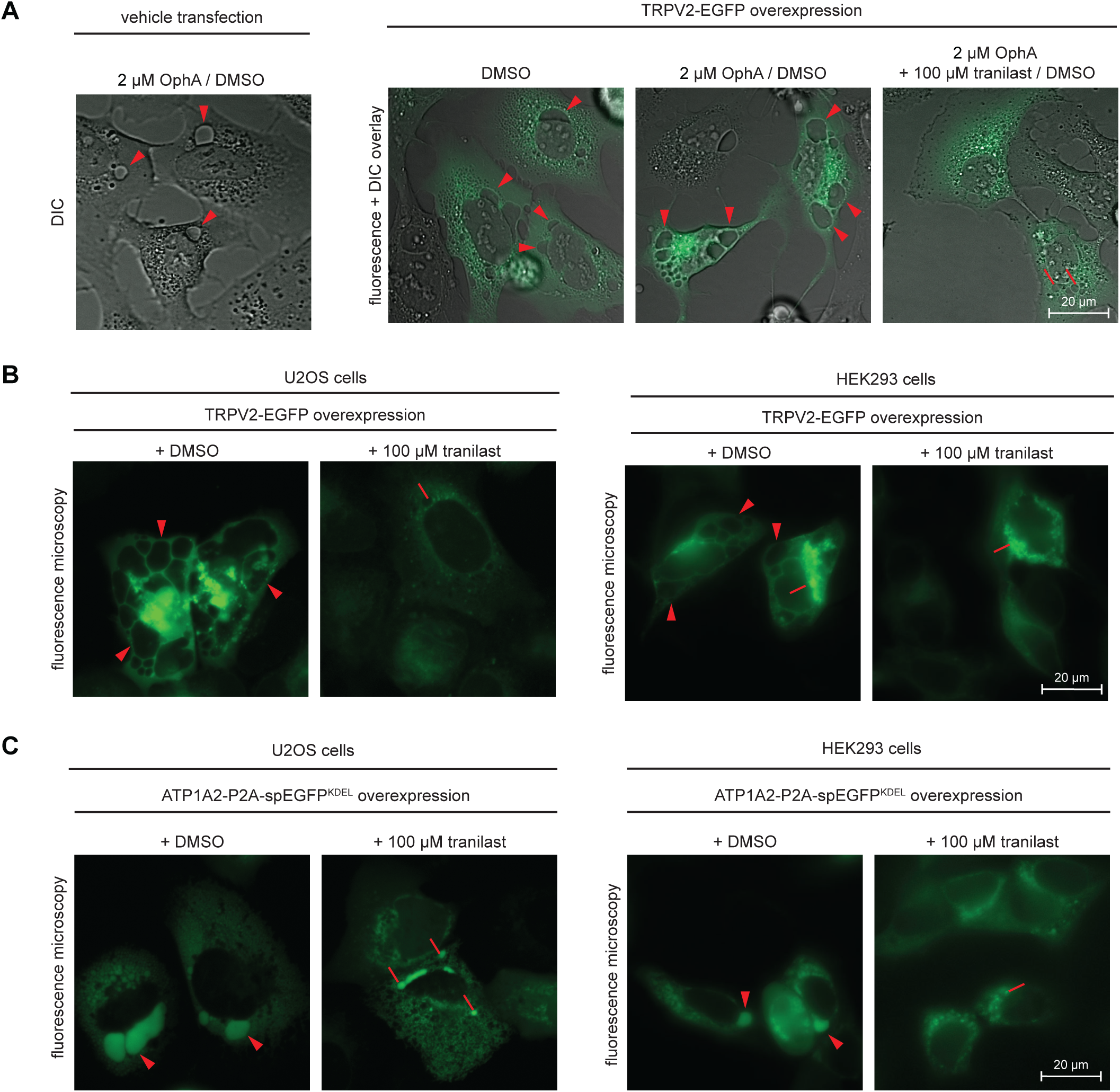
ER/PNS dilation phenotypes caused by TRPV2 or ATP1A2 overexpression can be blocked by tranilast. (A) Overexpression of TRPV2 fused to a C-terminal EGFP reporter causes vacuolation in the absence of OphA, and when combined with OphA treatment exacerbates vacuolation in a manner that can be blocked by tranilast. (B) The TRPV2 overexpression-induced vacuolation is not an idiosyncrasy of U2OS cells but can also be observed in HEK293 cells and can be prevented in both cell types by addition of 100 µM tranilast to the cell culture medium. (C) ER/PNS dilations induced by ATP1A2 and visualized by concomitant expression of spEGFP can be observed in U2OS cells and HEK293 cells and can blocked by tranilast. Red arrowheads and lines indicate PNS dilations and smaller ER vacuoles, respectively.

To explore if this phenotype is an idiosyncrasy of the U2OS cell model or can also be induced in other cells, we compared side-by-side the result of overexpressing TRPV2^EGFP^ in HEK293 cells. In this second human cell model, we observed both the formation of vacuoles and their rescue with tranilast (**Fig 6B**). We then investigated whether tranilast could also reverse the ER and PNS dilations induced by ATP1A2 overexpression in these *in vitro* paradigms and found that it effectively restored these alterations (**Fig 6C**). Overall, these results suggest that ER and PNS dilations in several paradigms depend on a common mechanism involving the disruption or overload of the ER-resident TRPV2 channel. This disruption leads to cellular Ca^2+^ influx and an imbalance in Ca^2+^ homeostasis across the ER lipid bilayer. If this imbalance persists, it may cause osmotic inflation of the subcellular compartment, unless TRPV2 is kept from translocating to the plasma membrane.

### OphA induced ER and PNS dilation is accompanied by an increase in cytosolic and ER calcium levels that can be blocked by a cell permeable calcium chelator

To determine if OphA poisoning is associated with Ca^2+^ influx into the cytosol and ER, we repeated the OphA treatment of U2OS cells in the presence of fluorescent calcium sensors. When deploying the cytosolic calcium sensor Fluo-3, we observed an increase in cytosolic calcium levels 3 hours after OphA addition (**Fig 7A**). Because Fluo-3 is hindered from entering the ER and PNS, we repeated the analysis with the Mag-Fluo-4 derivative of this reporter that has been shown to access this compartment. This experiment documented bright calcium fluorescence within the dilated ER/PNS compartments 3 hours after OphA addition.

**Fig. 7.**
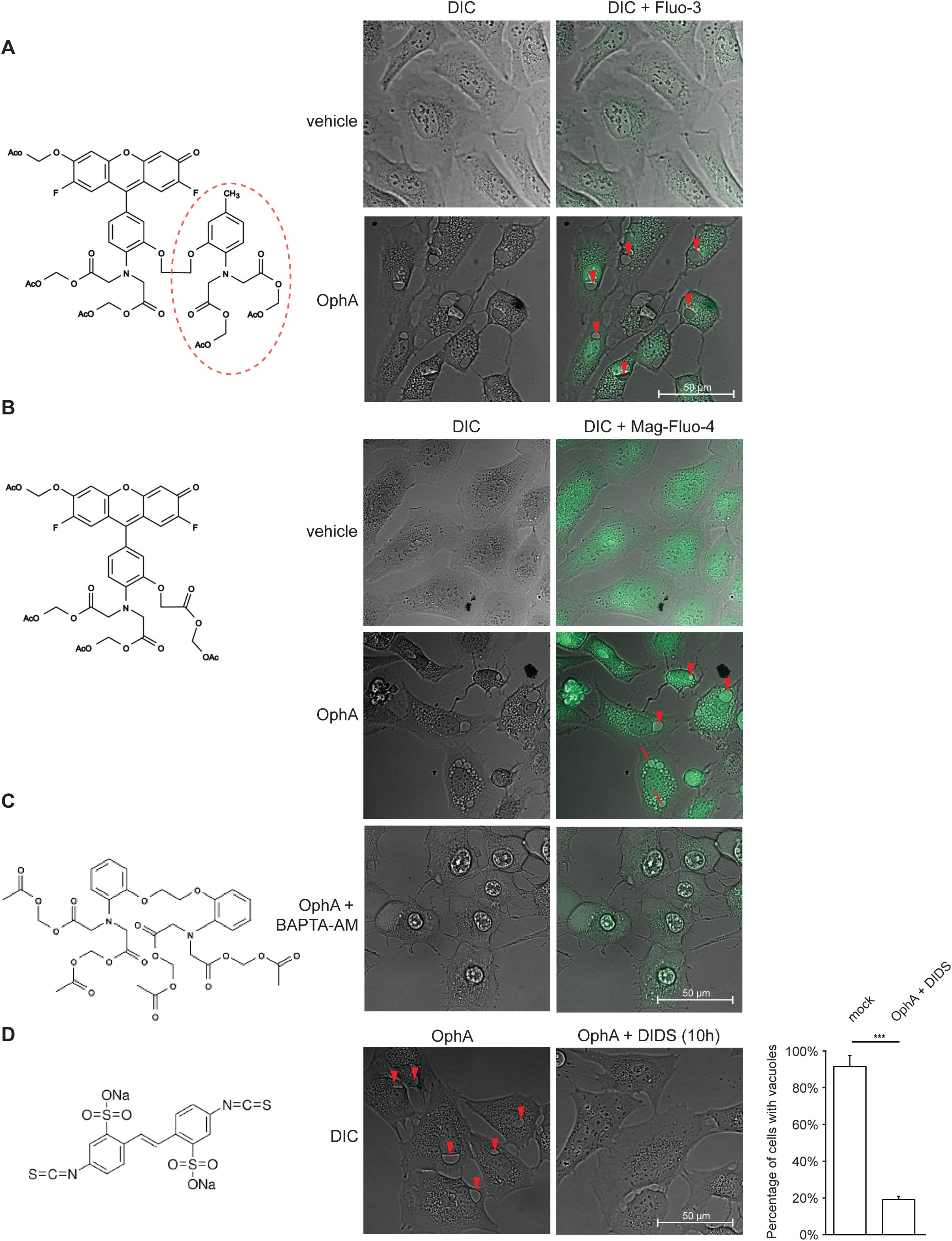
OphA induced ER/PNS dilation is accompanied by an increase in cytosolic and ER calcium levels that can be blocked by a cell-permeable calcium chelator. (A) OphA addition to U2OS cells leads to an increase in cytosolic calcium that can be visualized with the calcium reporter Fluo-3. (B) The ER penetrant calcium reporter Mag-Fluo-4 revealed robust calcium levels in the OphA-induced ER/PNS dilations. (C) Addition of the calcium chelator BAPTA-AM to cell culture medium of cells exposed to OphA prevented the increase in intracellular calcium levels and the formation of ER/PNS dilations. (D) U2OS cells exposed to OphA were prevented from developing ER/PNS dilations by the addition of DIDS, an agonist of ryanodine receptors that is understood to increase the opening probability of these channels leading to increased Ca2^+^ leakage from the ER into the cytosol. The graph to the right depicts the approximately 80% reduction in the percentage of cells exhibiting vacuoles in the presence of DIDS.

To determine if these apparent increases in Ca^2+^ influx represent mere bystander effects or are causative for the ER and PNS dilations, we conducted an OphA treatment in the presence of the cell-permeable calcium chelator BAPTA-AM and observed that its presence blocked the OphA induced swelling of the ER and PNS (**Fig 7C**). Finally, we wondered if the phenotype can be rescued by the addition of diisothiocyanostilbene-2′,2′-di*-*sulfonic acid (DIDS), a compound that has long been known to non-reversibly activate ER membrane-resident ryanodine receptor Ca^2+^ channels by increasing their open probability ^57–61^. As we had anticipated, this reagent was able to revert the ER and PNS dilation phenotype when added at known effective concentration exceeding 100 µM to cells that had been exposed to OphA (**Fig 7D**).

### PNS dilation induced by OphA can be blocked by heterologous expression of an N-terminal TRPV2 construct, responds in a complex manner to direct TRPV2 inhibitors, and is accompanied with CHOP induction and reduction in PIKFYVE levels

To further validate a possible role of TRPV2 in the ER and PNS dilation phenotype, we cloned an N-terminal TRPV2 construct (amino acids 1-387) fused C-terminally to the EGFP reporter (N-term-TRPV2-EGFP). The TRPV2 component in this fused construct comprised the cytoplasmic ankyrin repeat domain, which had been shown to interfere with TRPV2 cell surface localization ^55^ (**Fig 8A**). We then transfected U2OS cells with either the full-length TRPV2-EGFP construct alone or both full-length TRPV2-EGFP and the N-term-TRPV2-EGFP construct. When the full-length TRPV2-EGFP construct was overexpressed alone, we observed the expected dilations of the ER and PNS. When co-expressed with N-term-TRPV2-EGFP, this vacuolation phenotype was almost entirely rescued, except for the presence of the occasional vacuole seen in a small number of cells (**Fig 8B**).

**Fig. 8.**
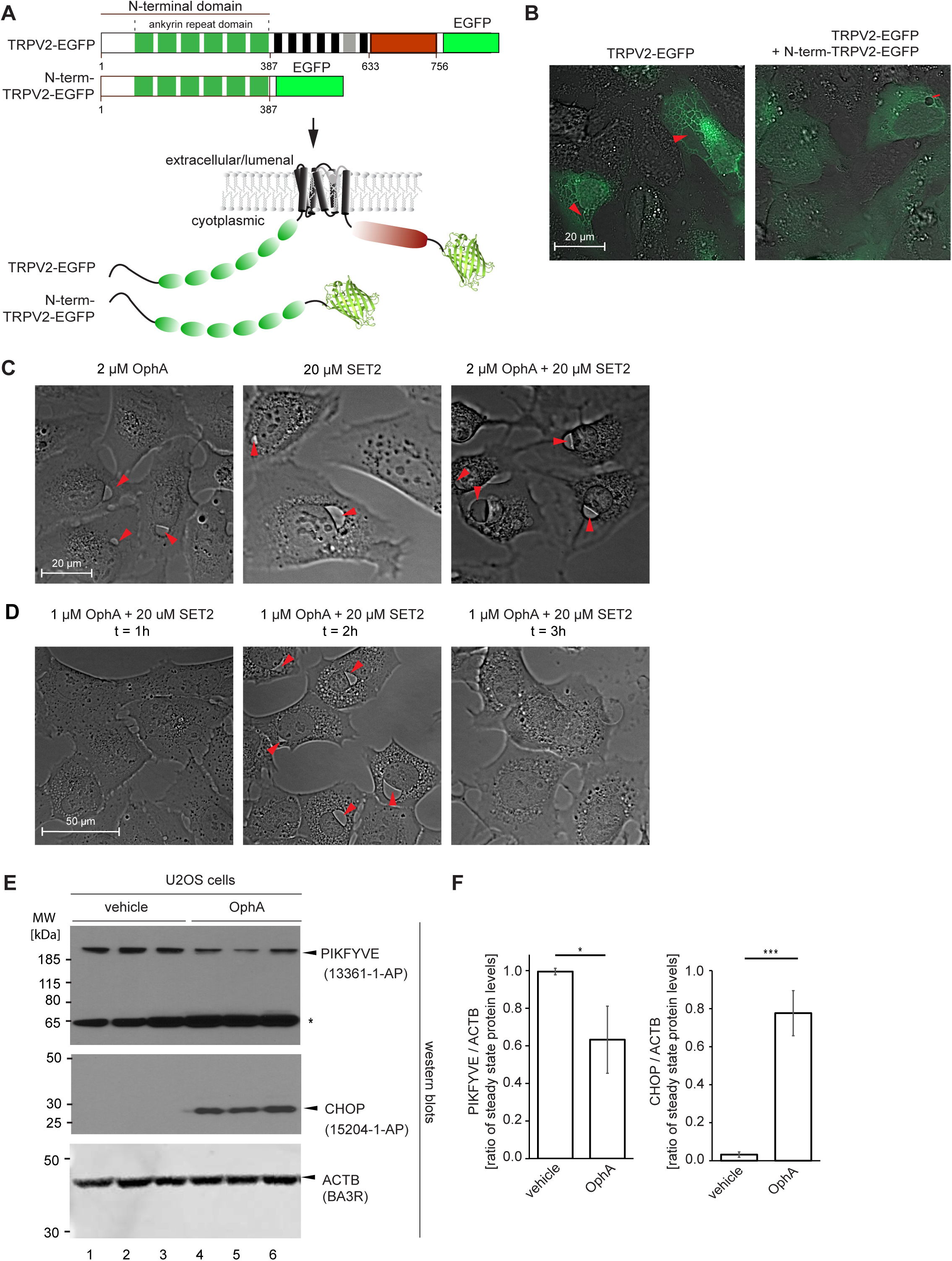
The PNS dilation phenotype can be blocked by overexpression of N-terminal TRPV2 or exposure to SET2 and is accompanied by CHOP upregulation and reduction in PIKFYVE levels. (A) Cartoon depicting domain organization and membrane topology of TRPV2 next to an N-terminal construct derived from it that comprises only the ankyrin repeat domain. Both expression constructs were C-terminally fused to EGFP. (B) Overexpression of TRPV2-EGFP causes a profound ER/PNS dilation phenotype that can be rescued by the co-expression of N- term-TRPV2-EGFP. (C) Exposure of U2O2 cells to OphA or the TRPV2-selective antagonist, SET2, causes U2OS cells to acquire PNS dilations. Concomitant exposure to both reagents induced a severe phenotype that, in addition to severe PNS dilation, was marked by increased cell shrinkage and early cell death. (D) Time-series of U2OS cells treated with both SET2 and OphA, the latter at a lower concentration of 1 µM, led to PNS dilation within 2 h, with reversal to normal cellular morphology 3 h after treatment onset. (E) Cells exposed to 2 µM OphA exhibit a reduction in steady-state PIKFYVE protein levels and an increase in CHOP protein levels. The asterisk points to an anti-PIKFYVE antibody cross-reactive band, whose identity was not pursued. (F) Graph documenting significant reduction in the ratio of PIKFYVE and ACTB protein levels in OphA-treated cells.

Next, we exposed U2OS cells to SET2, a selective TRPV2 antagonist that does not cross-react with other members of the TRPV family and is known to bind directly to its target ^62^. Microscopy analyses conducted two hours after adding 20 µM SET2 to the cell culture medium revealed that this antagonist induces a phenotypically similar ER and PNS dilation to that observed with 2 µM OphA. When both reagents were added concomitantly at these respective concentrations, the vacuolation phenotype was exacerbated, which was apparent by relatively larger PNS dilations and more severely shrunk nuclei (**Fig 8C**). Interestingly, when OphA was added alongside 20 µM SET2 to the cell culture medium at the lower concentration of 1 µM OphA, we saw the same phenotype forming after 2 h—albeit it at a slightly less severe level—yet microscopy analyses undertaken after 3 h revealed the U2OS to have reverted to their normal morphology without PNS dilation (**Fig 8D**). Such a phenotype reversal has not been observed in cells treated with OphA alone. Attempts to repeat the SET2-dependent observation led to mixed results, with the reversal being observed occasionally but not reliably. Because this observation can advance mechanistic interpretations (see discussion), we chose to document it here once we were able to capture it in live cell microscopy (see **S3 Video**). Taken together, these results suggested that OphA and SET2 act differently in this experimental paradigm despite both being able to induce the ER and PNS dilation phenotype. These observations align with a model in which SET2 initially induces ER and PNS dilation by antagonizing the calcium leak activity of ER-resident TRPV2. However, it may later rescue the dilation phenotype by blocking TRPV2- dependent Ca^2+^ influx through the plasma membrane when the cells have responded to ER folding stress with overexpression of the TRPV2 channel and its increasing translocation to the plasma membrane.

To begin to characterize the ER stress molecularly, we next added vehicle solution or OphA to U2OS cells, then harvested cellular extracts 2 hours later and analyzed them with an antibody directed against C/EBP homologous protein (CHOP). Mirroring prior results by others ^44^, we observed that steady-state CHOP protein levels robustly increased in OphA treated cells.

Next, we investigated steady state levels of PIKFYVE, a lipid kinase best known for its contribution to endolysosomal fission and fusion biology, whose functional ablation can lead to a pronounced vacuolation phenotype. We became interested in whether OphA treatment affects the levels of this lipid kinase because prion-infected brains were reported to exhibit reduced steady-state levels of PIKFYVE in a manner that appeared to be triggered by ER stress ^4^. These observations led the authors of this prior work to propose that vacuoles associated with spongiform degeneration in prion diseases are of endolysosomal origin. However, a definite determination of the nature of intracellular neuronal vacuoles—characteristic of the spongiform degeneration observed in prion-infected brains—remains elusive. We investigated if OphA-induced ER and PNS vacuolation is linked to a reduction in steady-state PIKFYVE levels and found them decreased by approximately 40% in the absence of endolysosomal vacuolation (**Fig 8A**).

### Direct evidence for ER/PNS dilation contributing to vacuolation in prion-infected mice

Finally, we revisited our original question: whether the type of abnormal vacuolation, now understood to represent ER and PNS dilation, contributes to vacuolation in prion-infected mice. To this end, we intracerebrally inoculated two cohorts of three 60-day-old C57Bl6 wild-type mice. One cohort received brain extracts from an uninfected mouse (control), while the other was inoculated with brain extracts from a mouse that had succumbed to prion disease following infection with Rocky Mountain Laboratory (RML) prions. 90 days later, we retro-orbitally administered rAAV vectors whose payload coded for spEGFP^KDEL^ into each mouse. To achieve efficient uptake across the blood brain barrier (BBB), the payload was encapsulated in the 9P31 capsid ^63^, known to dock to endothelial carbonic anhydrase (CAR4) for efficient CNS transduction ^64, 65^. At ∼160 days postinoculation (dpi), when the prion-inoculated mice exhibited the expected signs of late-stage prion disease, all mice were sacrificed, brain slices cryo-sectioned, and analyzed by fluorescence microscopy (**Fig 9A**).

**Fig. 9.**
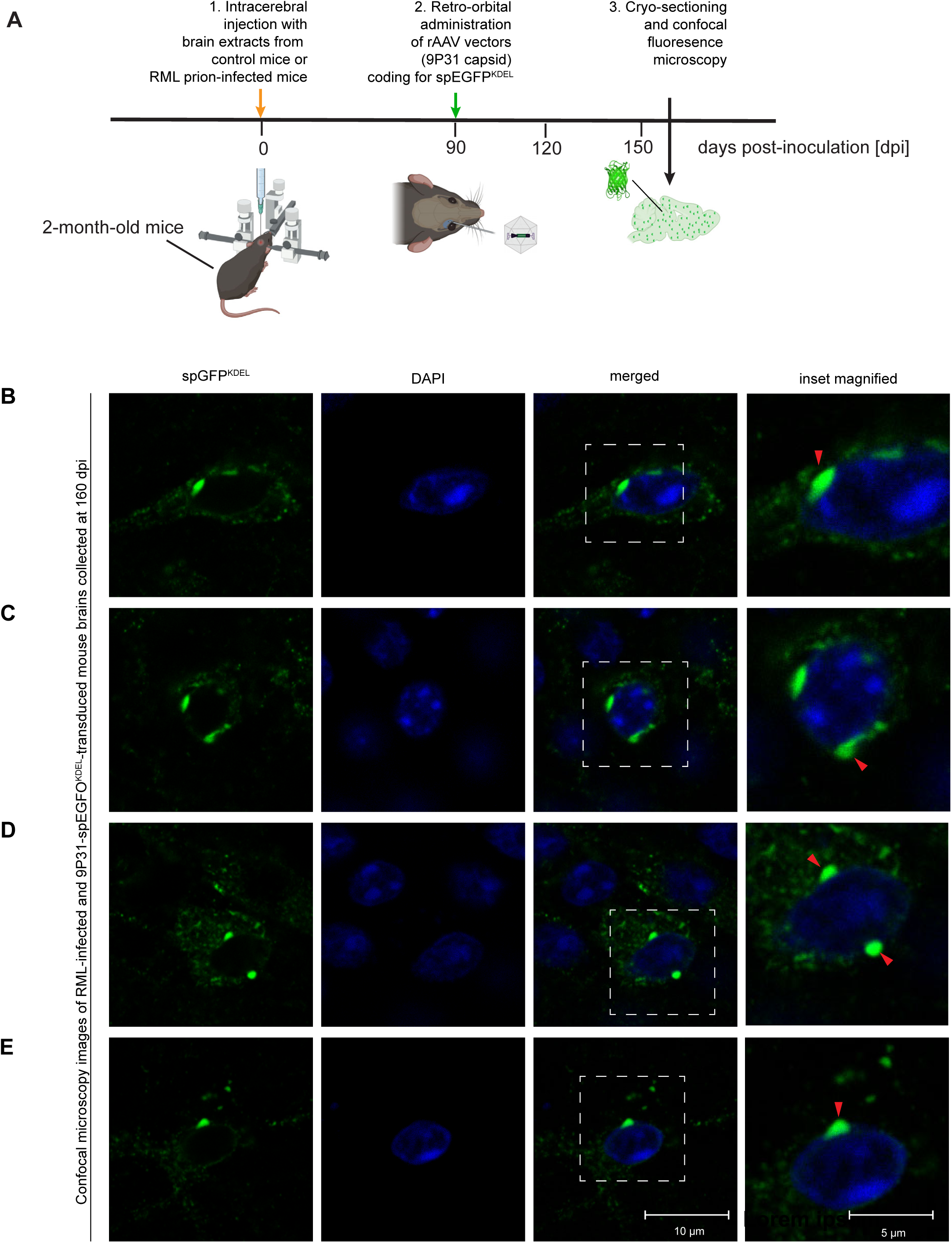
Prion infection of mice causes PNS dilations that can be positively identified in brain cryosections following spEGFP-KDEL transduction. (A) Schematic of experimental design. (B-E) Examples of confocal microscopy images of PNS dilations observed in late-stage RML prion-infected mice. The ER and PNS dilations fluoresce green because the mice had been transduced with rAAV vectors encapsulated in the 9P31 capsid with a payload coding for spEGFP^KDEL^.

In preparation of these analyses, we had investigated which kind of treatments can be inflicted on U2OS cells without losing the spEGFP^KDEL^ fluorescence signal within OphA-induced PNS dilations. These experiments revealed that the spEGFP^KDEL^ fluorescence within large PNS dilations is more susceptible to formaldehyde or glutaraldehyde exposure than the same signal observed associated with regular ER structures: Whereas the latter survived extended formaldehyde crosslinking, the spEGFP^KDEL^ signal within PNS dilations was mostly disappearing within 30-minutes of crosslinking with 1% formaldehyde at 4 °C (**S2 Fig A, B**). Moreover, although PNS dilations survived immersion in PBS or 30% sucrose (**S2 Fig C, D**), their spEGFP^KDEL^ signal disappeared during equilibration in a 1:1 (v/v) mix of 30% sucrose and O.C.T. freezing medium, a step described in several cryo-preservation protocols (**S2 Fig E**). Accordingly, in preparation of cryo-sectioning the mouse brains, we perfused them with PBS, then immersed them in 30% sucrose, followed by rapid equilibration and freezing in O.C.T. To minimize the introduction of cutting artefacts of this non-fixed tissue, we employed a film-backing method, also known as the Kawamoto method, which greatly stabilizes tissue sections ^37^.

The subsequent confocal fluorescence analyses of cryo-sections from prion-infected brains revealed several instances of PNS dilations. These dilations were identified based on a) their invariable juxtaposition to nuclei, b) smooth and relatively wide green fluorescence, c) and persistent fluorescence across several adjacent layers within confocal Z-stacks. Although readily identifiable within neurons in deeper brain structures, these PNS dilations were never abundant, i.e., at most revealing three such structures within one field of view (comprising approximately one hundred brain cells) (**Fig 9 B-E**). In all instances captured, the sizes of these PNS dilations were smaller than the sizes of adjacent nuclei, and as such were smaller than the biggest PNS dilations we had observed in cell-based studies. To date, we could not resolve if a higher fragility of larger PNS dilations placed an upper limit on the size at which these structures can survive the tissue handling steps we had decided on. No such PNS dilations were found in age-matched healthy control mice. These experiments provided direct *in vivo* evidence for the manifestation of ER/PNS dilations in prion diseases, raising the prospect that this phenotype contributes to the spongiform degeneration that represents one of the pathological hallmarks of prion diseases.

## DISCUSSION

The objective of this study was to investigate if our recent discovery of the close spatial relationship between PrP^C^ and NKA can advance understanding of how neuronal vacuolations develop in prion diseases. This objective led us on a protracted experimental journey that brought to the fore several pertinent insights: 1) Neuronal vacuolation can be induced in mice whose Atp1a1 subunit has been humanized by changing two amino acids that govern CG binding; 2) Overexpression of NKA α subunits induces vacuoles in a subset of human cells; 3) The dilated compartments in these paradigms are cisternae of the ER and the PNS, whose osmotic swelling can also be induced by the overexpression of a subset of other transmembrane proteins or by exposure to certain toxins known to induce ER stress; 4) The apparent turgor observed in ER and PNS swellings is caused by increases in calcium levels in these cisternae and is independent of sodium or potassium ion exchange activities of NKAs. 5) The ER and PNS swellings can be induced by antagonizing the ER calcium release function of the TRPV2 channel and can be prevented by chelating calcium, by hindering the translocation of TRPV2 to the plasma membrane, or by expressing an N-terminal TRPV2 truncation construct. 6) PNS dilations can also be documented in prion-infected animals.

To our knowledge, this study is the first to link the vacuoles observed in prion diseases to an ER and PNS dilation phenotype, which can be induced by various cellular insults, including CG poisoning, overexpression of certain multi-spanning transmembrane proteins, and exposure to specific toxins. Additionally, this study may be the first to successfully fill vacuoles observed in prion-infected mouse brains *in vivo* with a fluorescent marker protein.

Because the PNS represents the largest continuous ER cisterna in most cells, profound ER swelling is most readily observed as a PNS dilation. When encountered, this phenotype is characterized by the presence of translucent PNS vacuoles that can sometimes occupy most of a cell’s cytoplasm. The absence of electron dense structures in these vacuoles is inconsistent with protein densities being the main cause of osmotic imbalances. Instead, these vacuoles appear to form due to an internal turgor caused by a small osmotically active compound, most likely an ion.

This study is not the first to seek a mechanistic explanation for the formation of large ER and PNS vacuoles. For instance, a prior report suggested that OphA exposure of cells disrupts the activity of the big conductance Ca^2+^-activated K^+^ channel (BKCa), thereby affecting the cellular potassium ion homeostasis ^45^. The authors also noted increases in cytosolic calcium levels and ascribed them to compensatory ER mechanisms that may involve IP3 or ryanodine receptors. Our own data place stronger emphasis on cellular calcium influx and overload within the ER and PNS being the cause for its osmotic swelling based on our observations that 1) the cell membrane-permeable calcium chelator BAPTA-AM could prevent the distention of the ER/PNS, 2) the TRPV2 channel, which we deem central to this event in the OphA paradigm, is a calcium release channel, and 3) we observed a transient translocation of TRPV2 from the ER to the plasma membrane upon OphA treatment that preceded the forming of the PNS dilations. We wonder if a separate study, which documented that BKCa channel-mediated potassium extrusion works synergistically with the TRPV2 dependent calcium influx ^66^ may reconcile data from this study and the prior report.

Moreover, it had previously been reported that the compound DIDS can block profound swelling of the ER and PNS (and mitochondria) induced by a prenylated flavonoid, termed morusin ^67^. The authors concluded that this outcome relied on inhibition of the voltage-dependent anion channel (VDAC) by DIDS. An alternative scenario may be considered, i.e., rather than ascribing the effect of DIDS to an inhibitory role toward VDAC, we favor the interpretation— consistent with data by others ^57, 59, 60^—of DIDS opening calcium release channels in the ER, including the ryanodine receptor, thereby preventing the build-up of calcium in this compartment.

Cells react to mild non-toxic CG levels with internalization of NKAs, along with the raft domains they are inserted in, leading to the co-internalization and degradation of the prion protein ^68–70^. In our previous work, we observed repeatedly that as we increased CG concentrations toward toxic levels, cells ramped up NKA expression, presumably to replace inhibited NKAs at the cell surface ^68^. This cellular response has the potential to induce ER stress if it fails to replenish functional NKAs at the cell surface. If the process is unsuccessful, the accumulation of dysfunctional NKAs could overwhelm the ER, triggering stress. One way in which PrP^Sc^ might lead to a PNS dilation phenotype therefore would be if PrP^Sc^ aggregates poison raft domains and interfere with raft-resident NKAs, triggering a futile cellular response that aims to replace functional raft domains in a subset of brain cells in late-stage prion disease (**Fig 10**).

**Fig. 10.**
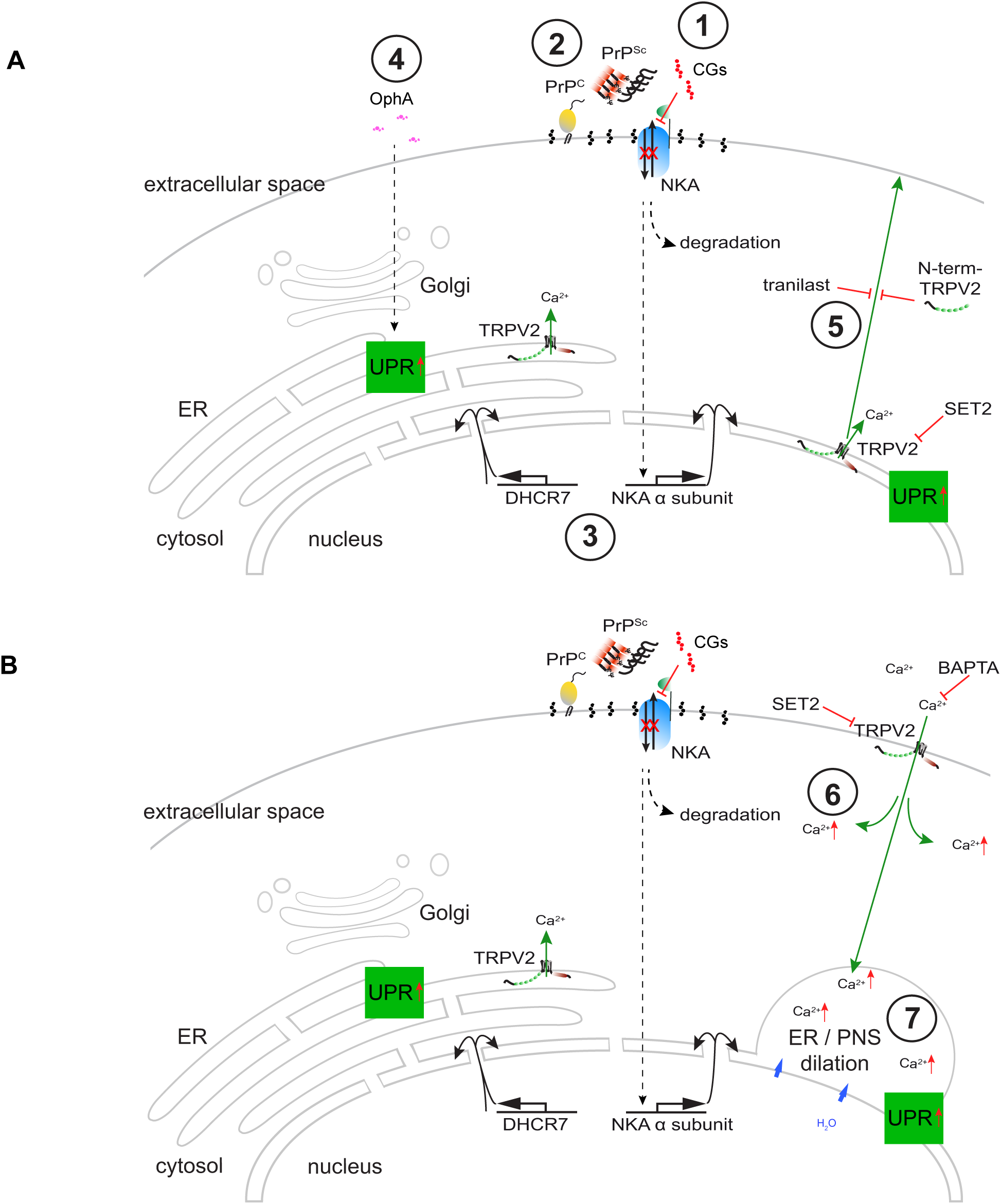
Cartoon depicting model of molecular events underlying ER and PNS dilation. (A) Model summarizing insults that can lead to ER stress, induction of the UPR, and PNS dilation. 1. Acute CG poisoning causes cells to internalize and degrade NKAs. This induces cells to increase expression of replacement NKAs, which may eventually overwhelm the ER. 2. Accumulation of PrP^Sc^ may poison raft membrane domains, causing cells to increase expression of raft-resident transmembrane proteins, including NKA subunits, which may again overwhelm the ER. 3. Mere overexpression of certain transmembrane proteins causes phenotypically similar ER stress. 4. Exposure of cells to certain xenotoxins, e.g., OphA, blocks progression of proteins along the secretory pathway, leading to ER stress. 5. Cells appear to counteract this type of ER stress by increasing expression TRPV2 and possibly translocating some of it to the plasma membrane. This can be blocked by tranilast or by overexpression of a truncated N-term-TRPV2 construct, which may compete with endogenous TRPV2 for factors involved in this biology. (B) 6. Plasma membrane inserted TRPV2 increases ER calcium levels by facilitating cellular calcium entry into the cell and by not fulfilling its ER calcium leak channel activity. This biology can be blocked by chelating calcium with BAPTA-AM or by with the TRPV2-specific antagonist SET2. 7. Elevated ER calcium levels cause osmotic swelling of the ER, which may facilitate protein folding by reducing crowding and providing calcium to many proteins, including GRP78, that require calcium as a co-factor when assisting in rescuing the cell from ER stress. This process may initially remain reversal but leads to cell death if swelling becomes excessive.

Several natural toxins that induce the PNS dilation phenotype, including OphA and CsA, may similarly stall the progression of proteins along the secretory pathway and are therefore also bound to cause ER stress and induce the UPR ^44^. The observation that the membranes expand in these paradigms suggests ongoing biosynthesis of ER membranes. We therefore propose these phenotypes to represent cases of UPR activation in which the eIF2α phosphorylation-dependent slowing of protein synthesis is partially reversed by the selective translation of a subset of proteins that is governed by ATF4 and CHOP activation ^8, 71^. This cellular response is reminiscent of another ER phenotype that leads to an increase in ER membranes, known as ER hypertrophy. This phenotype was initially described in cells exposed to phenobarbital. It has been proposed that these cells attempt to outrun their demise by driving the massive production of drug metabolizing enzymes ^72^. Interestingly, this phenotype can also be induced by the overexpression of certain membrane proteins, including proteins involved in sterol biosynthesis, not unlike what we and others have observed in DHCR7 overexpression paradigms ^73^. These similarities indicate that it may be rewarding to compare the molecular underpinnings of these programs more closely.

A strength and limitation of this study is its relatively broad scope, spanning several ER and PNS dilation phenotypes and cellular insults that ranged from acute toxin poisoning to protein overexpression to prion disease. This design element was introduced to overcome certain limitations, e.g., a reality whereby large vacuoles that can easily be captured by light microscopy, are not routinely observed in cell models used in prion infection studies, although vacuoles have been documented to form in some paradigms ^4, 74, 75^. The obvious caveat of this approach is that it had to rely on an assumption that distinct stressors, which cause ER and PNS dilation, do so through common molecular underpinnings. Although this simplifying hypothesis is not far-fetched, it will almost certainly require revision and refinement. For instance, whereas vacuolation observed upon OphA exposure or overexpression of NKA α subunits appears to center on TRPV2, other PNS dilation phenotypes may be linked to alternative modes of cellular Ca^2+^ influx. A hint of further complexity is indicated by recent studies, which proposed that TRPM3 and 8 channels can also be influenced by OphA ^76^.

Another limitation of this study is its sole focus on the ER and PNS dilation, a deliberate choice because it was the most striking morphological change we observed. As the number of reports documenting similar phenotypes—now most often referred to as paraptosis—has grown to just over one hundred studies ^34, 77^, several authors have shown that ER and PNS dilations are often accompanied by swelling of mitochondria and are associated with increases in mitochondrial calcium ^78–80^. Moreover, there already is considerable data on the subset of UPR markers whose levels change in various ER dilation paradigms. This prior work has most consistently revealed CHOP levels to be increased, a data point we confirmed in this study. A more complex molecular fingerprint of the UPR governing paraptosis is coming into focus that includes the expected concomitant upregulation of ATF4, phosphorylation of c-jun N-terminal kinase-1 (JNK-1) ^81, 82^, as well as increases in BiP and IRE1α ^44, 67, 83^. Finally, proteasome inhibition—detectable by an increase in ubiquitination—seems to be an additional molecular hallmark of this phenotype ^34, 77^. Straightforward experiments that explore the presence of these markers in prion-infected cells and tissues may offer rewarding angles for validating and adding granularity to the data reported here.

Although we provide direct *in vivo* evidence for PNS dilations in prion diseases, we are hesitant to speculate on the significance of this finding for understanding the spongiform degeneration phenotype without additional corroborating data. The notion that spongiform degeneration might be based on ER and PNS dilations—although not dominant in the prion literature—is not new, perhaps most forcefully proposed based on neuropathological analyses of marmosets infected with Creutzfeldt-Jakob disease (CJD) or Gerstmann-Sträussler-Scheinker (GSS) prions ^28^. More recently, the presence of dilated ER and PNS cisternae was also noted in an ultrastructural investigation of lesions in sheep afflicted with natural scrapie ^75^. Our close inspection of published and unpublished images of spongiform degeneration in various prion disease paradigms revealed many depictions of large vacuoles adjacent to nuclei (for examples see images in ^28, 84^). In these images, the respective nuclei are often shown to be dented on the side where the vacuoles abut and exhibit various degrees of nuclear shrinkage of the kind we observed in OphA-treated cells. A more systematic analysis of the neurohistopathological record, including early ultrastructural analyses, may be instructive. Such an analysis may be enhanced by a comparison of endolysosomal vacuoles observed in mouse brains deficient for Pikfyve activity ^85, 86^. For now, we showed that steady-state level reductions in PIKFYVE can also be observed in human cells exposed to OphA, which are marked by profound ER and PNS dilations—not endolysosomal vacuoles.

Preferably, direct labeling of spongiform degeneration vacuoles will further clarify the relative contribution of intracellular compartments to the manifestation of this disease phenotype. As more than hundred years of uncertainty regarding the origins of spongiform degeneration attest, this objective is not trivial and conventional neuropathological methods, including formaldehyde crosslinking, followed by paraffin-embedding, and H&E staining, i.e., the dominant tissue processing pipeline for documenting spongiform degeneration, is unlikely to reveal the answer. Previous attempts to depict spongiform degeneration in cryo-sectioned brain sections were described as problematic as it was found to be difficult to discern vacuoles with this sample preparation method ^87^. In this study, filling the ER with spEGFP^KDEL^, avoidance of formaldehyde crosslinking and O.C.T. equilibration, followed by cutting relatively thick 40 µM brain sections (to minimize physical disruption of vacuoles) led us to detect the green fluorescence-filled PNS dilations in prion-infected mice. However, these PNS dilations were never abundant and smaller than the largest ones that we observed in other *in vitro* paradigms of ER and PNS dilation. OphA poisoning led cells to burst and die within a few hours. Given this relatively short timeline, the detection of a few PNS dilation within a field of view may not so much indicate this event to be rare but too fleeting for the detection of large numbers of spEGFP^KDEL^-filled PNS dilations. Refinement of methods will be required to settle this question and possibly capture larger spEGFP^KDEL^-filled PNS dilations, if they exist.

## ACKNOWLEDGMENTS

The authors are grateful to the late Jerry B. Lingrel (University of Cincinnati) for generously allowing the use of *Atp1a1^s/s^*knock-in mice for this study. We also thank Nickolai O. Dulin (University of Chicago) for shipping a breeding pair of this mouse line to Toronto.

## SUPPLEMENTARY INFORMATION

**S1 Fig. Suppression of ATP1A2 initially appeared to generate the ER/PNS dilation phenotype, an observation was revealed to depend on the rare occurrence of partially digested siRNAs causing the upregulation of a target gene through mRNA stabilization.**

(A) Treatment of U2OS cells with siRNAs targeting ATP1A1 or ATP1A2 validated a reduction in steady-state NKA α1 levels but not α2 levels. (B) Although pool A of ATP1A2-directed siRNAs did not achieve the reduction in steady-state levels of its protein target, it induced the ER/PNS dilation phenotype. (C) RNAse A digestion of pool A of ATP1A2-directed siRNAs blocked its induction of ER/PNS dilation, indicating this phenotype to be dependent on the presence of RNA in this pool. (D) Side-by-side comparisons of pool A with newly synthesized individual siRNAs present within this pool or additional pools A and B from the same versus a different vendor document reduced levels of ATP1A2-directed siRNA in pool A. (E) RT-qPCR-based analyses of ATP1A1 and ATP1A2 transcripts in U2OS cells that had been exposed to pools A-C revealed pool A to increase steady-state mRNA levels, as opposed to decreasing them, whereas pools A and B led to the expected suppression of ATP1A1 mRNA levels.

**S2 Video. Time-lapse video of OphA-induced PNS dilations in U2OS cells.**

U2OS cells were treated with 2 µM OphA, followed by live cell microscopy capture of cells beginning at 1.5 h after the addition of OphA. The 18 s long time-lapse video captures a 5-hour time window, with images taken every 5 minutes. At the onset of the video several cells in the periphery have already developed PNS dilations. Other cells, seen in the center of the field of view, are just beginning to develop this phenotype. The nuclei of cells exhibiting this phenotype appear to condense as the PNS dilations increase in size. As the phenotype becomes more pronounced, the nuclei of the respective cells are deformed at sites of contact to the PNS dilations. Smaller ER cisternae can also be observed to expand. Eventually, several cells detach from the cell culture dish. Other cells seem to disintegrate into smaller vesicles, with some of these vesicles merging and eventually giving rise to a large bubble surrounding the ‘ghost’ cell. A link to the video will be made available during peer review.

**S3 Video. Time-lapse video of SET2-dependent reversal of PNS dilation.**

1 µM OphA and 20 µM of the TRPV2 inhibitor SET2 were concomitantly added to the cell culture medium of U2OS cells. The time-lapse sequence captured a ∼30-minute time window in a 13 s video, with images taken at a rate of 1 image per minute. The video begins 2 h after the addition of the reagents, when several cells exhibited profound PNS dilations. During the sequence several cells can be seen to gradually reverse the PNS dilation phenotype. A link to the video will be made available during peer review.

**S4 Fig. Fixation of cells or freezing in OCT cryo-embedding medium removes fluorescence signal from PNS dilations.**

U2OS cells were transfected with the spEGFP^KDEL^ expression plasmid, followed by addition of 2 µM OphA to the cell culture medium after 48 h. Prior to capturing fluorescence microscopy and differential inference contrast (DIC) images at 2 h after OphA addition, the cell culture medium was removed and replaced with solutions as indicated for the durations shown. (A) Exposure to PBS did not impact detection of spEGFP^KDEL^ fluorescence in PNS dilations (shown with red arrowheads). (B) Mild formaldehyde (FA) crosslinking (0.5%, 15 min) also preserved spEGFP^KDEL^ fluorescence within PNS dilations. (C) Stronger fixation in the presence of 2% FA and for a duration of 15 minutes retained the translucent shape of PNS dilations (shown with white arrowheads) but was not compatible with the subsequent detection of spEGFP^KDEL^ fluorescence within PNS dilations. (D) Mild crosslinking with glutaraldehyde also canceled spEGFP^KDEL^ fluorescence within PNS dilations. (E) 30-minute embedding of cells in 30% sucrose in PBS preserved spEGFP^KDEL^ fluorescence in PNS dilations. (F) Pre-equilibration in 1:1 (v/v) of 30% sucrose in PBS : O. C. T. freezing medium preserved the translucent PNS dilations but precluded the subsequent direct detection of spEGFP^KDEL^ fluorescence within PNS dilations. Note that all treatments shown preserved the green fluorescence of spEGFP^KDEL^ in smaller ER cisternae.

## Abbreviations

BBB: blood brain barrier
CG: cardiac glycoside
CHOP: C/EBP homologous protein
CJD: Creutzfeldt-Jakob disease
CsA: cyclosporin A
CWD: chronic wasting disease
DHCR7: 7-dehydrocholesterol reductase
DIDS: 4,4′-diisothiocyanatostilbene-2,2′-disulfonic acid disodium salt hydrate
EGFP: enhanced green fluorescent protein
ER: endoplasmic reticulum
ITR: inverted terminal repeat
NCX: sodium calcium exchanger
NKA: Na^+^,K^+^-ATPase
OphA: ophiobolin A
PNS: perinuclear space
PrP: prion protein
PrP^C^: cellular prion protein
PrP^Sc^: scrapie prion protein
rAAV: recombinant adeno-associated virus
RML: Rocky Mountain Laboratory
SET2: N-(Furan-2-ylmethyl)-3-((4-(N7-methyl-N′-propylamino)-6-(trifluoromethyl)- pyrimidin-2-yl)thio)-propanamide
TRPV2: transient receptor potential vanilloid 2
TSE: transmissible spongiform encephalopathy
UPR: unfolded protein response.

