## Supplementary figures and images for "Transient receptor potential vanilloid channel 2 contributes to multi-modal endoplasmic reticulum and perinuclear space dilations that can also be observed in prion-infected mice"

### Supplemental Figures

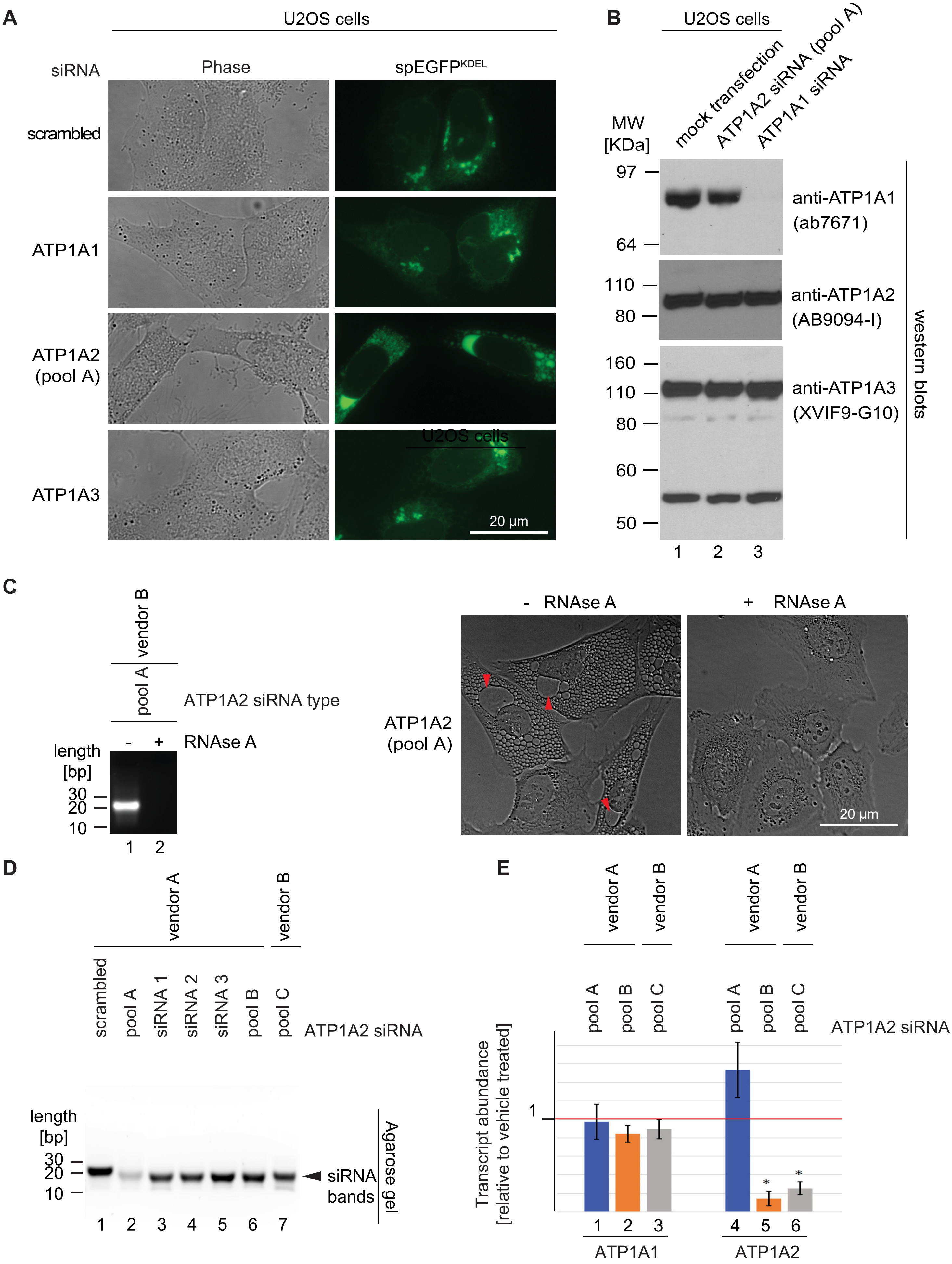

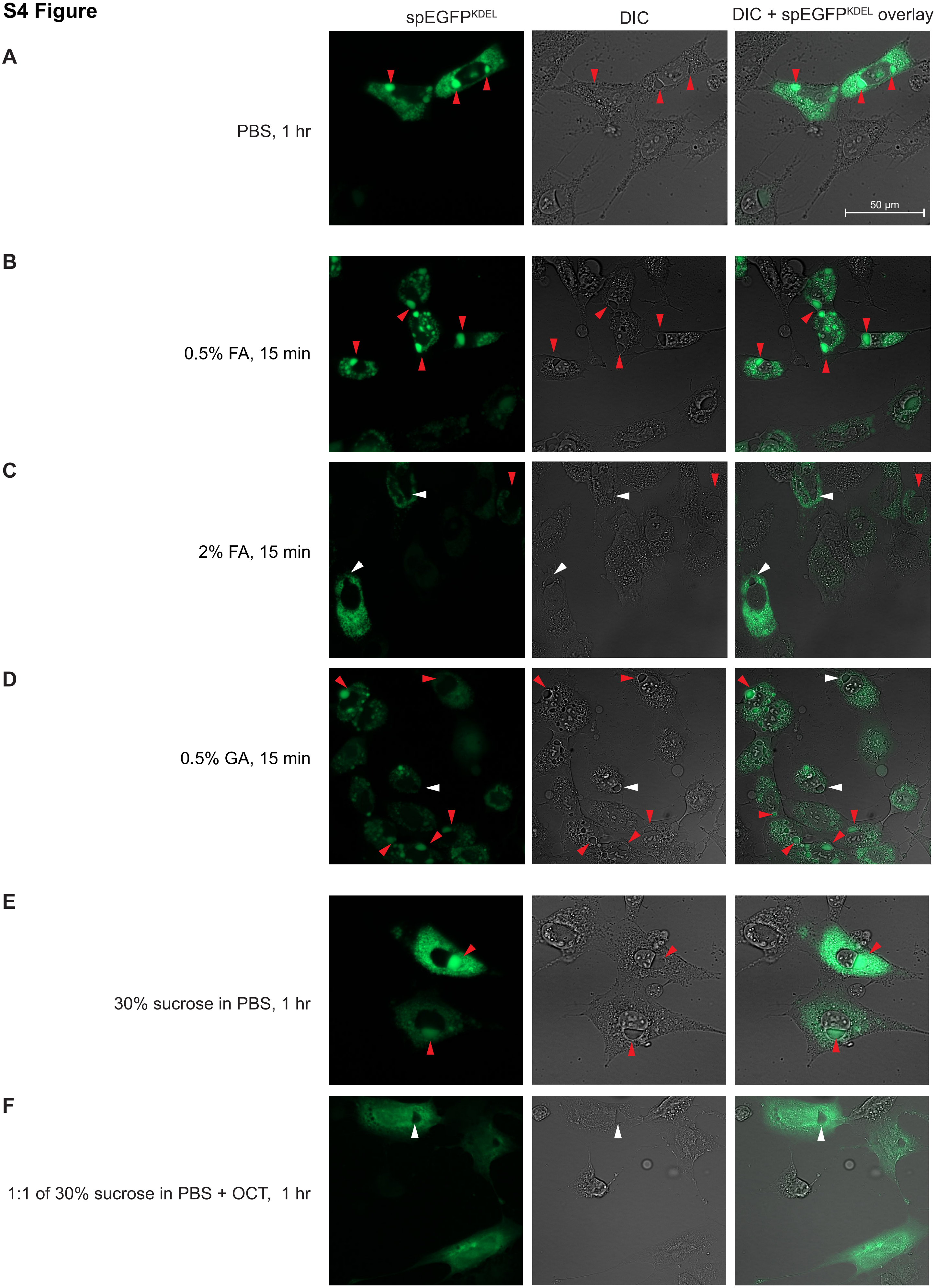
